# Visible traits demonstrate that crispant founder mice can be used for phenotypic assessment

**DOI:** 10.1101/2025.06.27.661753

**Authors:** Rebekah Tillotson, Marina Gertsenstein, Li-Hsin Chang, Julie Ruston, Fernando Bellido Molías, Lauri G. Lintott, Christine Taylor, Philippe Gautier, Lauryl M. J. Nutter, Monica J. Justice

## Abstract

Genes can be knocked out in model organisms by introducing a single guide RNA and Cas9 into one cell zygotes. Recently, the zebrafish and *Xenopus* communities have employed this method in genetic screening pipelines that assess phenotypes in founders (F0), referred to as “crispants”. In contrast, phenotyping crispant mice has been avoided as results are believed to be confounded by genetic mosaicism, requiring that only established mouse lines undergo phenotypic assessment. Here, we targeted seven genes associated with visible recessive phenotypes. We observed the expected null phenotype in up to 100% founders per gene. Crucially, we achieved 100% editing efficiency in all but two animals. Genetic mosaicism was common, but did not confound an animal’s phenotype when comprised of mutations that all disrupted the targeted gene. Mosaicism included short in-frame mutations, but these were sufficient to disrupt function of five genes. Several founders were compound heterozygotes carrying a null and a non-null allele (short in-frame mutation or late truncation), enabling functional assessment of the non-null allele to dissect protein function. Our results set the stage for using crispant founders for initial phenotypic assessment in genetic screening, before selecting candidates for further study. This will dramatically reduce animal numbers.

## Introduction

Phenotypic analysis of single and multi-gene mutants is critical for assessing the function of individual genes, redundancy between related genes, and genetic interactions. While molecular and cellular phenotypes resulting from genetic disruption in monocellular organisms and cultured cells can be informative, whole animals are required to identify affected tissues throughout development and assess more complex phenotypes. Model organisms used by geneticists include *Drosophila melanogaster* (fruit fly), *Caenorhabditis elegans* (nematode worm), *Xenopus laevis* (African clawed frog) and *Danio rerio* (zebrafish), but the mouse is the “workhorse” for understanding mammalian genetics and modeling human disease.

Experimental techniques can be divided into two categories: forward genetics (phenotype-driven) and reverse genetics (gene-driven)^1^. In mice, forward genetics began with the collection of animals with visible traits by 19^th^ century “mouse fanciers”. These traits occurred naturally in wild populations or arose spontaneously in laboratory colonies. Subsequently, the mutation rate was enhanced by mutagenising the male germline with radiation (primarily X-rays) or chemicals (primarily N-ethyl-N-nitrosourea, ENU) or by “gene-trapping” using transposons (Sleeping Beauty or PiggyBac). Of these, the “supermutagen” ENU is most frequently used to screen for phenotypic traits^2^. ENU is a DNA alkylating agent that creates small lesions, mostly single base pair changes, creating hypomorphic/hypermorphic/neomorphic/dominant-negative/null/neutral alleles. Dominant traits (e.g. development of cataracts) can be detected in G1 offspring derived from the mutagenised males. Recessive traits (e.g. embryonic lethality) require further breeding of G2 females with G1 males to achieve homozygosity of *de novo* mutations in G3 offspring^3,4^. ENU mutagenesis has also been used to screen for dominant suppressors of genetic diseases (thrombocytopenia, Rett syndrome and Factor V Leiden), revealing key disease pathways and druggable targets^5–8^. The greatest challenge in these screens is identifying the causative mutation(s). ENU mutagenesis is random and G1 pups carry ∼40-70 intragenic lesions (and many more intergenic ones). The vast majority of these have no/little phenotypic consequence and mutation(s) underlying observed phenotypes must be determined in pedigrees derived from the G1 animals by linkage analysis or by a combination of whole exome sequencing (WES) and genotyping of G3 animals. If G1s cannot be bred due to fertility or health problems, then identification of causative mutations is not possible. Overall, ENU screening involves several challenges and huge numbers of animals (especially for recessive traits), highlighting a clear need for a more efficient and cost-effective method.

The sequencing and annotation of model organism genomes has revolutionised genetic studies. Of ∼20,000 protein-coding genes in the human genome, 80-90% are conserved in mice^9^. Reverse genetics approaches predominantly involve knocking out individual genes to produce null alleles. Such studies have been carried out at a small scale by individual laboratories on their genes of interest, and at a large scale by the International Mammalian Phenotyping Consortium (IMPC). The IMPC’s aim is to identify the function of every protein-coding gene in the mouse genome by producing a knock-out mouse line for each^10^. Initially, mouse lines were created by gene targeting in cultured embryonic stem (ES) cells, which were then used to produce chimeras comprising both wild-type and mutant ES cells. If the mutant cells contributed to the germline, mutant mouse lines could be established upon breeding these chimeras. Over the last decade, CRISPR/Cas9 technology has been adopted to improve the efficiency of gene editing in ES cells^11^, and – more importantly – to perform gene targeting directly in zygotes^12–14^. This has accelerated mouse model production and enabled mutations to be introduced onto different strain backgrounds (circumventing the need for backcrossing). There are several ways to deliver the single guide RNA (sgRNA) and Cas9 nuclease to zygotes, with the best results obtained when recombinant sgRNA and Cas9 protein are combined to form a ribonucleoprotein (RNP), which is introduced into zygotes via electroporation^14^. Null alleles can be produced by excising a critical exon with sgRNAs targeting either side; or by creating insertions/deletions (indels) within a critical exon using one sgRNA and relying on imperfect repair via non-homologous end joining (NHEJ) or microhomology-mediated end joining (MMEJ). Founders are bred and quality control assays are performed on N1 animals to verify disruption of the targeted gene before mouse lines are established and analysed^15^.

CRISPR/Cas9 technology has also been adopted for forward genetic screening of cellular phenotypes in both cultured cells and *in vivo* in model organisms (including mice) by introducing Cas9 with a genome-wide or bespoke library of sgRNAs at a low multiplicity of infection such that each cell carries one sgRNA and, as a result, its target gene is disrupted^16–18^. To assess whole animal phenotypes, recent studies using zebrafish and *Xenopus* have performed *in vivo* genetic screens for phenotypes of interest by introducing RNPs into zygotes and phenotyping the resulting F0 founder animals^19–23^. These founders are named “crispants”, a portmanteau of CRISPR and mutant. Crispants are genetically mosaic, but they display the null phenotype associated with the targeted gene due to high levels of indel formation. The resulting mutations can be identified by sequencing the sgRNA target sites in the founders. As both copies of the targeted gene are disrupted, recessive phenotypes can be assessed.

The mouse community remains sceptical about whether phenotypic assessment of F0 crispant mice can produce reliable results, with initial studies reporting phenotypic variability due to genetic mosaicism^24^. Here, we target seven genes associated with recessive visible traits such as coat colour and demonstrate that up to 100% of crispant mice per gene displayed the expected null phenotype despite genetic mosaicism. Our results were due to very high levels of Cas9- mediated double-stranded breaks and the resulting production of loss-of-function mutations. We then asked whether it was possible to target two genes per zygote as this would be desirable when assessing genetic redundancy or genetic interactions. Unfortunately, we found that this decreased the viability to a level that made it not worthwhile. Genes would therefore need to be targeted sequentially in zygotes to introduce secondary mutations. We obtained a relatively high proportion of non-mosaic founders that were compound heterozygous for two null alleles or one null allele and one short in-frame mutation. When compound heterozygous founders carrying a short in-frame mutation displayed the recessive loss-of-function phenotype, this mutation pin- pointed a critical functional protein domain. Overall, our findings demonstrate the validity of phenotypic analysis of crispant mice.

## Results

### Crispant mice display recessive coat colour phenotypes

Phenotypic assessment of mouse crispant founders is generally avoided due to fears that genetic mosaicism would invalidate results. To determine whether phenotypic assessment of founders is biologically informative, we selected genes that when mutated present with recessive visible phenotypes. Mice with altered appearance first became popular among mouse fanciers and there are now over 300 “named” alleles – named after their phenotypes^25^. Most of the genes responsible and the causative mutations have now been identified. We chose genes for this study that have a recessive phenotype, no impact on viability or health, and a phenotype visible on a C57BL/6J background. We mimicked previously published null alleles (exon deletion or nonsense mutation) by targeting affected exons with a single sgRNA, introducing insertions/deletions (indels) from the repair of Cas9-induced double stranded breaks via NHEJ/MMEJ. We used strict off-target criteria^13^ whereby all sgRNAs had a least three mismatches to unintended targets, at least one of which was in the 11 bp “seed” region adjacent to the PAM, and all sgRNAs had a predicted frameshift frequency >67% (assessed using inDelphi^26^). We first selected two coat colour genes, as any phenotypic mosaicism would be clearly visible, allowing us to distinguish between a full, partial or absent knock-out phenotype and link this to the mutations carried by each founder.

Mutations in *Tyrp1* (tyrosinase related protein 1) result in brown coat colour on a nonagouti (a/a) background. Although the original “brown” allele, *Tyrp1^b^*, is a SNP (c.329G>A) resulting in the p.C110Y missense mutation^27^, many other historical “brown” mouse lines contain large deletions at this locus. We therefore decided to knock out *Tyrp1* by targeting a constitutive, translated exon located in the first half of the gene and with a length indivisible by three. The gene is composed of 8 exons, with alternative splicing affecting the length of exons 1 and 8. The open reading frame starts in exon 2 and ends in exon 8. We targeted exon 3 (Figure 1A; Table S1), previously deleted by the KOMP project in ES cells^28^ (*Tyrp1^tm1a(EUCOMM)Wts^*^i^). We electroporated 19 mouse zygotes with RNP and produced four founder animals (21% viability), all of which had a fully brown coat colour (Figure 1B; Table 1).

**Figure 1:**
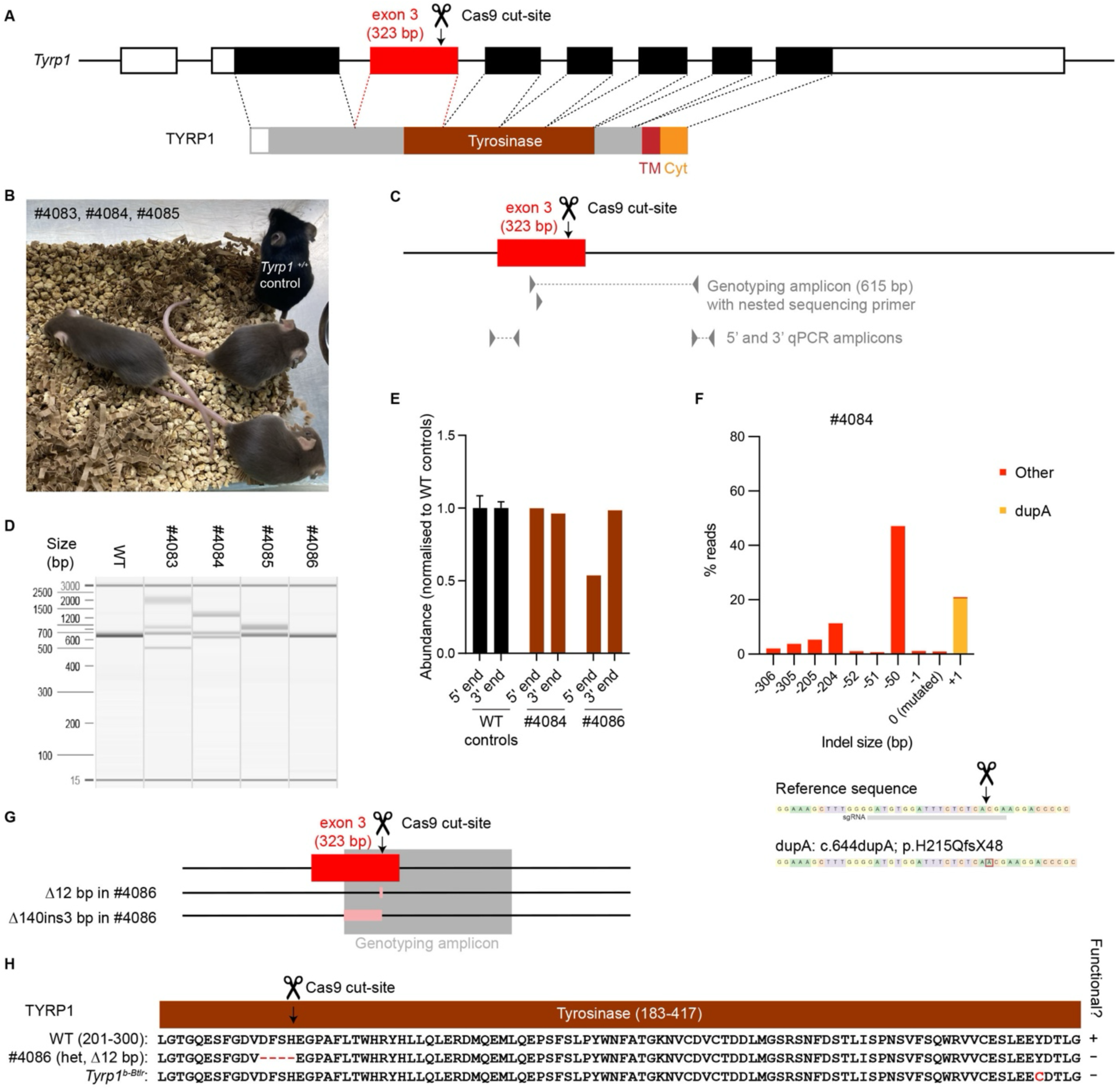
Crispant mice with mutated *Tyrp1* display the expected recessive brown coat colour phenotype. A. Schematic of the *Tyrp1* gene with the predominant TYRP1 protein isoform below (exons and protein domains drawn to scale). The position of the Cas9 cut site is indicated by the scissors and arrow. Key: 5’ and 3’ UTRs: white; open reading frame (ORF): black; targeted exon: red; excised propeptide: white; tyrosinase domain: brown; transmembrane domain “TM”: red; cytoplasmic domain “Cyt”: orange; and uncharacterized protein regions: grey. B. Photograph of founders with a *Tyrp1^+/+^*control. C. Schematic of exon 3 of the *Tyrp1* gene showing the location of the Cas9 cut site, genotyping and qPCR primers (drawn to scale). D. PCR amplification around the targeted site in all four *Typr1* founders, beside a wild-type reference (615 bp). E. qPCR quantification around the 5’ and 3’ genotyping primer annealing sites in founders #4084 and #4086, normalised to three wild-type controls (mean ± SEM). F. DAS analysis of #4084 (percentage abundance of indels present in >0.5% reads are shown). G. Schematic of the two alleles in the compound heterozygous founder, #4086 (drawn to scale, deleted regions shown in pink). H. Schematic of the protein sequence resulting from the in-frame deletion in #4086 (c.636_647del; p. D212_H215del) and the published *Tyrp1^b-Btlr^* allele.

**Table 1:** Overview of zygote EP experiments.

| **Gene** | **Viability** | **Phenotype** |
| --- | --- | --- |
| *Tyrp1* | 4/19 = 21.1% | Brown: 4/4 = 100% |
| *Slc45a2* | 21/39 = 53.8% | Underwhite (light grey), full: 2/21 = 9.5%<br>Mosaic ("chimera-like"): 6/21 = 28.6%<br>Black (as WT): 13/21 = 61.9% |
| *Fgf5* | 10/26 = 38.5% | Angora (long fur): 10/10 = 100% |
| *Tgm3* | 8/26 = 30.8% | Wellhaarig (curly coat and whiskers): 8/8 = 100% |
| *Bmp5* | 12/27 = 44.4% | Short ear: 12/12 = 100% |
| *Pax1* | 5/20 = 25% | Undulated (shorter kinky tail): 5/5 = 100% |
| *Adamts20* | 7/26 = 26.9% | Belted (white belt), full: 3/7 = 42.9%<br>Belted, partial: 3/7 = 42.9%<br>Black (as WT): 1/7 = 14.3% |
| *Tyrp1* + *Tgm3* | 1/23 = 4.3% | Wellhaarig + brown: 1/1 = 100% |
| *Tyrp1* + *Bmp5* | 1/23 = 4.3% | Short ear + brown: 1/1 = 100% |
| *Bmp5* + *Fgf5* | 0/82 = 0% | N/A |
| *Adamts20* + *Tgm3* | 4/74 = 5.4% | Belted + Wellhaarig: 4/4 = 100% |
| *Tyrp1* + *EGFP* | 5/26 = 19.2% | EGFP negative + Brown: 4/5 = 80%<br>EGFP negative + Patchy brown: 1/5 = 20% |
| *Fgf5* + *EGFP* | 1/26 = 3.8% | EGFP negative + Angora: 1/1 = 100% |
| *Tgm3* + *EGFP* | 7/39 = 17.9%<br>(Batch 1: 3/26 = 11.5%;<br>Batch 2: 4/13 = 30.8%) | EGFP negative + Wellhaarig: 7/7 = 100% |
| *Bmp5* + *EGFP* | 4/26 = 15.4% | EGFP negative + Short ear: 4/4 = 100% |

PCR amplification of the targeted locus from tail-tip genomic DNA (gDNA) of three founders (#4083, #4084 and #4085) produced at least one band that differed in size from the wild-type reference, indicating the presence of large indels (Figure 1C-D). Sanger sequencing of the amplicons produced mixed chromatograms that we deconvoluted using DECODR^29^ and Synthego ICE^30^ software (Figure S1; Table S2). DECODR was better at detecting longer deletions, so this software was primarily used throughout this study.

Since the trait is recessive, we were surprised that DECODR detected 26.9% wild-type (unedited) sequence in founder #4084, suggesting editing occurred in 3/4 alleles at the 2-cell stage, leaving 1/4 alleles unedited. This would mean that 50% cells carry a functional allele, which is at odds with the absence of black fur in this animal. One possibility was that this founder carried larger “unseen” deletions that abolish one or both primer annealing sites, so the PCR amplicon represented only a subset of the total alleles and therefore overestimated the proportion of unedited sequence. Quantification by qPCR around the genotyping primer annealing sites revealed that this is not the case (Figure 1C,E). Deep amplicon sequencing (DAS) revealed that a similar proportion of reads (20.4%) contained a duplication of an adenine base (c.644dupA; p.H215QfsX48; Figure 1F). We therefore conclude that DECODR (and ICE) mis- assigned this sequence as unedited when it actually contains an out of frame (+1 bp) insertion, consistent with the fully brown coat phenotype.

Surprisingly, founder #4086 appeared to carry a single allele: a short in-frame (12 bp) deletion (c.636_647del), resulting in the loss of four amino acids (p.D212_H215del) (Figure 1G-H, S1; Table S2). It seemed unlikely that this founder would be homozygous for this allele, as the same repair event would had to have happened in both copies of *Tyrp1,* and this repair product was not predicted by inDelphi. qPCR analysis around the genotyping primer annealing sites found that gDNA abundance was decreased by ∼50% over the 5’ primer (Figure 1E), suggesting that this founder was heterozygous for a “unseen” deletion spanning the 5’ primer annealing site. To identify the “unseen” deletion in founder #4086, we adapted ELF-CLAMP (<u>E</u>nrichment of <u>L</u>ong DNA <u>F</u>ragments using <u>C</u>apture and <u>L</u>inear <u>Amp</u>lification)^31^ where regions of interest (up to 2 kb) are amplified from viewpoints using *in vitro* transcription followed by Nanopore long read RNA sequencing (Figure S2A). We detected a large deletion-insertion (Δ140insCTT) within exon 3, which would lead to a frameshift mutation: c.794_933delinsCTT; p.I169Lfs*49 (Figure 1G, S2B). This was confirmed by PCR amplification and Sanger sequencing. Although a 140 bp deletion at a single cut site is relatively large, there are reports of on-target CRISPR/Cas9- mediated deletions in mouse zygotes up to 600 bp^32^.

With the knowledge that both alleles must be disrupted to cause the brown phenotype, both alleles in this compound heterozygous founder (#4086) must be non-functional, indicating that the four amino acid deletion is sufficient to destroy TYRP1 protein function. However, a report that CRISPR/Cas9-mediated mutagenesis in cell lines can cause genome damage in cis with small *de novo* mutations^33^ raises the alternative possibility that undetected damage nearby may be responsible for disrupting *Typr1*. Using ELF-CLAMP, we sequenced the region flanking this mutation and did not detect any additional mutagenesis, ruling this out (Figure S2B). The 12 bp (four amino acid) deletion is located in the tyrosinase domain (residues 183-417), and therefore demonstrates the functional importance of this domain *in vivo*. Consistent with this, a chemically induced (ENU) allele with a missense mutation in the tyrosine domain, *Tyrp1^b-Btlr^*; c.887A>G; p.Y296C (Figure 1H), also confers a brown coat colour. Overall, all four founders reliably displayed the null phenotype because we achieved complete editing and all *de novo* mutations carried by these animals were loss-of-function alleles (including short in-frame indels).

We next targeted a second coat colour gene, *Slc45a2* (solute carrier family 45, member 2), which when mutated results in a light grey coat on a nonagouti background – named “underwhite”^34^.

The gene is composed of seven constitutive exons and the open reading frame spans them all (Figure 2A; Table S1). The *Slc45a2^uw^* allele contains a 7 bp deletion in exon 3, truncating the protein (c.773_779delCGTCGCT; p.S258CfsX44)^35^. As we were unable to identify an sgRNA in this exon that met our off-target criteria, we instead mimicked *Slc45a2^uw-10Btlr^*, which phenocopies *Slc45a2^uw^* mice and encodes a nonsense mutation in exon 1 (c.39T>A; p.Y13X).

**Figure 2:**
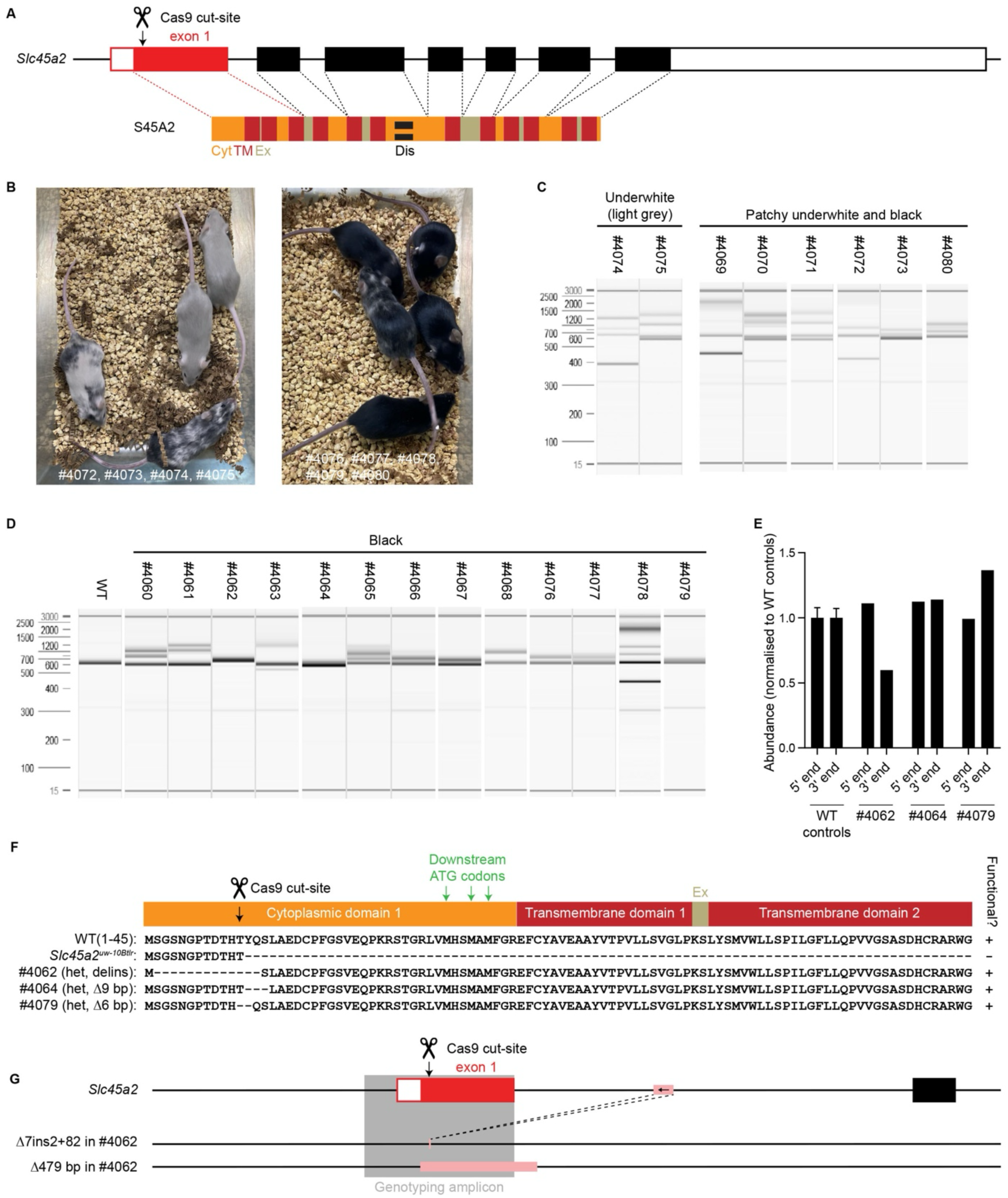
Crispant mice with mutated *Slc45a2* show phenotypic variability. A. Schematic of the *Slc45a2* gene with the S45A2 protein below (exons and protein domains drawn to scale). The position of the Cas9 cut site is indicated by the scissors and arrow. Key: 5’ and 3’ UTRs: white; open reading frame (ORF): black; targeted exon: red; transmembrane domains “TM”: red; cytoplasmic domains “Cyt”: orange; extracellular domains “Ex”: beige; and disordered “Dis”: black bands. B. Photographs of selected founders with a full light grey coat (left), patchy grey and black coat (left and right) and black coat (right). C-D. PCR amplification around the targeted site in light grey and patchy founders (C) and black founders (D), beside a wild-type reference (614 bp). E. qPCR quantification around the 5’ and 3’ genotyping primer annealing sites in compound heterozygous black founders (#4062, #4064 and #4079), normalised to three wild-type controls (mean ± SEM). F. Protein sequence resulting from the published truncated *Slc45a2^uw-10Btlr^* allele and in-frame indels in the compound heterozygous black founders. G. Schematic of the two alleles in the compound heterozygous founder with genome damage, #4062 (drawn to scale, deleted regions and the inserted region of intron 1 are shown in pink).

Electroporation of 39 zygotes resulted in 21 founders (53.8% viability; Table 1). Only two (9.5%) displayed the full underwhite phenotype, six (28.6%) had a patchy grey and black coat, and the remaining 13 (61.9%) appeared wild-type (Figure 2B). This result could not be explained by a lack of mutagenesis as no unedited sequence was detected in any founders (Figure 2C-D; Table S2).

We therefore asked whether we could determine the functional importance of the mutated N-terminal cytoplasmic domain from genotypic analysis. PCR amplification resulted in a single band for three phenotypically wild-type founders (#4062, #4064 and #4079) (Figure 2D).

Founders #4064 and #4079 were compound heterozygotes carrying two short deletions at a 1:1 ratio, one of which was in-frame: c.35_43delCCTATCAAT; p.T12T,Y13_S15del in #4064 and c.32_37delATACCT; p.H11H,T12_Y13del in #4079 (Figure 2E-F, S3; Table S2). We conclude that short in-frame deletions in the N-terminal cytoplasmic domain do not disrupt S45A2 protein function. Founder #4062 was a compound heterozygote carrying a complex deletion-insertion comprising a short deletion and an insertion including 82 bp from intron 1 (c.36_42delCTATCAAins[AA,385+647_385+566]), and a 479 bp deletion that was detected by ELF-CLAMP (c.1-1_385+93del) (Figure 2E,G, S3; Table S2). Since the underwhite phenotype is recessive, only one of these needs to be functional for the mouse to appear wild-type. The deletion-insertion provided an alternative start codon, predicted to enable the translation of a protein containing a short deletion, p.S2_Q14del (Figure 2F, S3). The finding that short-in frame mutations at the target site do not abolish S45A2 protein function explains the phenotypic variability in these founders.

### Crispant knock-out mice display recessive visible phenotypes

We selected four more genes linked to recessive visible phenotypes to knock out in crispant founders. Two genes affect the coat (ectoderm): *Fgf5* (fibroblast growth factor 5) and *Tgm3* (transglutaminase 3), associated with the named alleles “Angora” (long coat) and “Wellhaarig” (curly coat and whiskers), respectively^36,37^. The other two genes have roles in the mesoderm: *Pax1* (paired box 1) mutated in “undulated” (short wavy tail) and *Bmp5* (bone morphogenetic protein 5) mutated in “short ear” mice^38,39^. As with *Tyrp1* and *Slc45a2*, we took the gene structures into account and designed an sgRNA for each gene to mimic an existing knock-out allele^36,39–41^ (Figure 3A-D; Table S1). We electroporated 20-27 zygotes per gene and produced 5- 12 founders for each, giving a viability of 25-44% per gene. All founders displayed the expected recessive visible phenotypes (Figure 3E-H; Table 1, S2).

**Figure 3:**
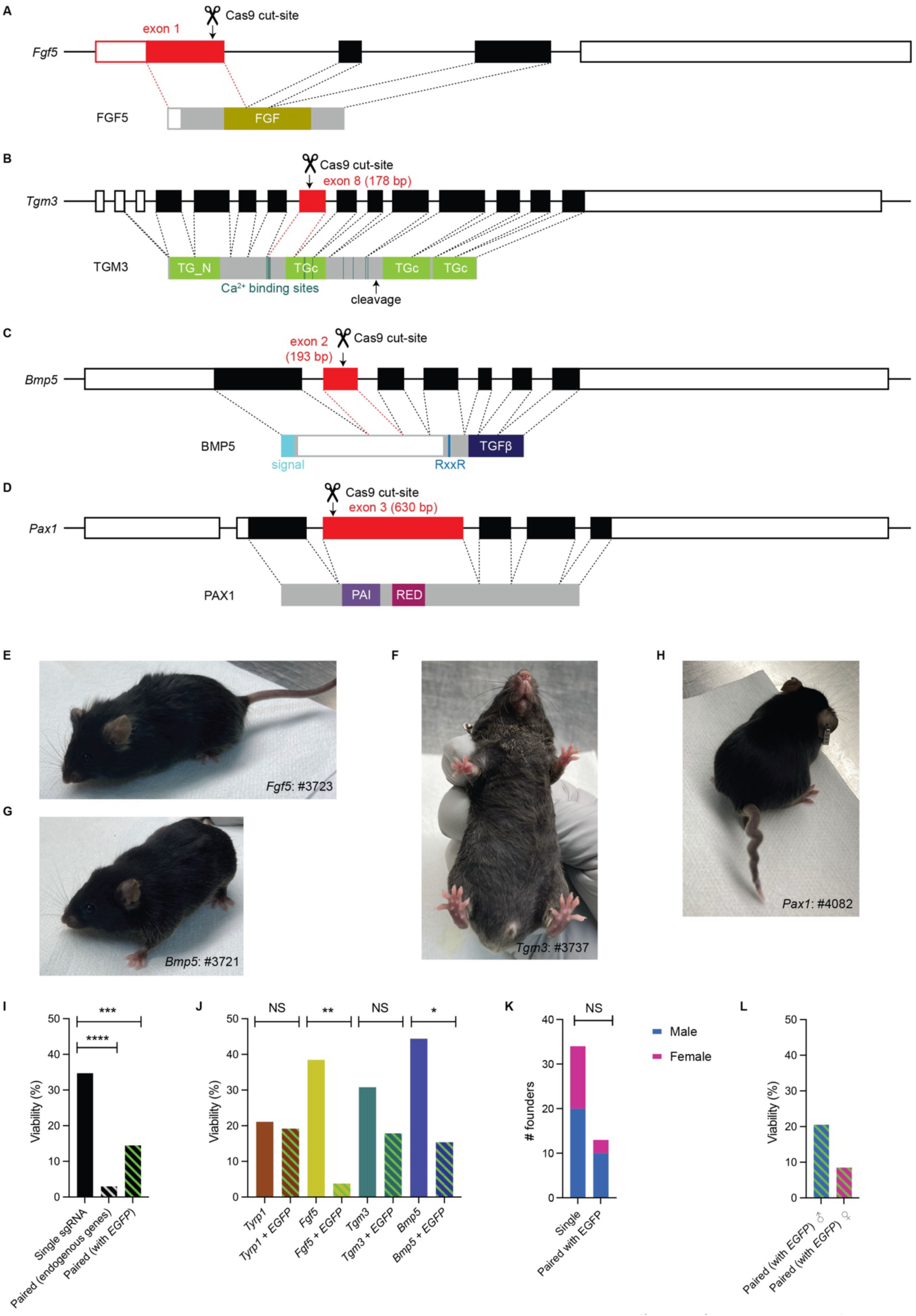
Crispant mice display recessive visible phenotypes. A-D. Schematic of the *Fgf5* (A), *Tgm3* (B), *Bmp5* (C) and *Pax1* (D) genes with their predominant/sole protein isoforms below (exons and protein domains drawn to scale). The position of the Cas9 cut site is indicated by the scissors and arrow. Key: 5’ and 3’ UTRs: white; open reading frame (ORF): black; targeted exon: red; excised propeptide: white; FGF domain: gold; N- and C-terminal Transglutaminase domains: green; calcium (Ca^2+^) binding sites: teal; signal peptide: cyan; RxxR motif: blue; transforming growth factor beta (TGFβ) like domain: navy; PAI-RED box domain first/second portion: purple/magenta and uncharacterised protein regions: grey. E-H. Photograph of one representative founder for each gene: *Fgf5* (E), *Tgm3* (F), *Bmp5* (G) and *Pax1* (H). I. Comparison of percentage viability when targeting one autosomal endogenous gene, two autosomal endogenous genes, or one autosomal endogenous gene in combination with X-linked *EGFP.* J. Comparison of percentage viability when targeting *Tyrp1*, *Fgf5*, *Tgm3*, and *Bmp5* alone or in combination with X-linked *EGFP.* K. Comparison of sex ratio of founders obtained when targeting one autosomal endogenous gene alone or in combination with X-linked *EGFP*. L. Comparison of estimated percentage viability when targeting one autosomal endogenous gene in combination with X-linked *EGFP* in male founders (three target sites per genome) and female founders (four target sites per genome). To calculate percentage viability in the endogenous gene + *EGFP* experiments, half of the 117 zygotes were assumed to be from each sex: males 12/58.5 = 20.5% and females 5/58.5 = 8.5%. I-L. Pairs of ratios were compared using Fisher’s Exact tests: **** P < 0.0001, *** P <0.001, ** P < 0.01, * P < 0.05 and NS (not significant) P > 0.05.

### Targeting two genes in crispants is possible but reduces viability

Targeting two genes at once in zygotes could be used to assess redundancy between related genes or to investigate genetic interactions, e.g. screening for modifiers of a chosen null phenotype. We tested four combinations of genes, ensuring pairs were on separate chromosomes to avoid large deletions. To avoid reagent toxicity, the total amount of RNP used was the same as for the single sgRNA experiments, with half the RNP containing each sgRNA. The viability was very low: 6/202 = 3.0% (**** P < 0.0001, Fisher’s Exact Test; Figure 3I; range per experiment: 0-5.4%; Table 1). The surviving founders, however, did display both expected recessive phenotypes (Figure S4).

Compared to experiments targeting single loci, the reduction in viability could be due to toxicity from the increased number of Cas9-induced breaks per cell and/or to synthetic lethality between the pairs of genes. To investigate this, we paired one sgRNA targeting an endogenous gene (*Tyrp1*/*Fgf5*/*Tgm3*/*Bmp5*) with an sgRNA targeting EGFP in transgenic zygotes carrying an EGFP transgene^42^. From a total of 117 zygotes, we recovered 17 pups at weaning (14.5%). This was significantly lower than when these four genes were targeted alone (*** P = 0.0007, Fisher’s Exact Test; Figure 3I; Table 1). When the viability for each gene was compared to the same gene combined with EGFP (Figure 3J), viability was significantly reduced for *Fgf5* (** P = 0.0048) and *Bmp5* (* P = 0.035). There was also a downward trend for *Tgm3*, but no difference for *Tyrp1* (which had the lowest viability when targeted alone). Since disrupting EGFP shouldn’t impact health, we conclude that the generation and repair of more Cas9-induced breaks reduced viability.

The transgene was X-linked, allowing us to compare the impact of adding of one extra target site in hemizygous males vs two extra sites in homozygous females. The high percentage of males in the paired sgRNA experiments (12/17 = 71%) was indicative of increased toxicity from four target sites, but this sex ratio wasn’t significantly different from that obtained in the single sgRNA experiments due to low founder numbers (Figure 3K). Assuming equal numbers of males and females in the 117 zygotes used in the paired sgRNA experiments, the viability of male and female founders was 20.5% and 8.5%, respectively (Figure 3L). The viability of female founders was higher than when two endogenous genes were targeted (8.5% vs 3.0%), despite the same number of target sites. This suggests that synthetic lethality was also a factor.

Overall, the more than four-fold decline in viability makes targeting two genes together less efficient than targeting a second gene in zygotes that are homozygous for mutations in a first gene, derived from a heterozygous intercross (Mendelian ratio: 25%).

All 17 founders obtained in the paired sgRNA experiments lacked EGFP fluorescence and displayed the full KO phenotype associated with the targeted endogenous gene, with mutagenesis at both loci (Table 1). The only exception was one *Tyrp1 + EGFP* founder, #4728, which had brown fur with black patches (Figure 4A, discussed later).

**Figure 4:**
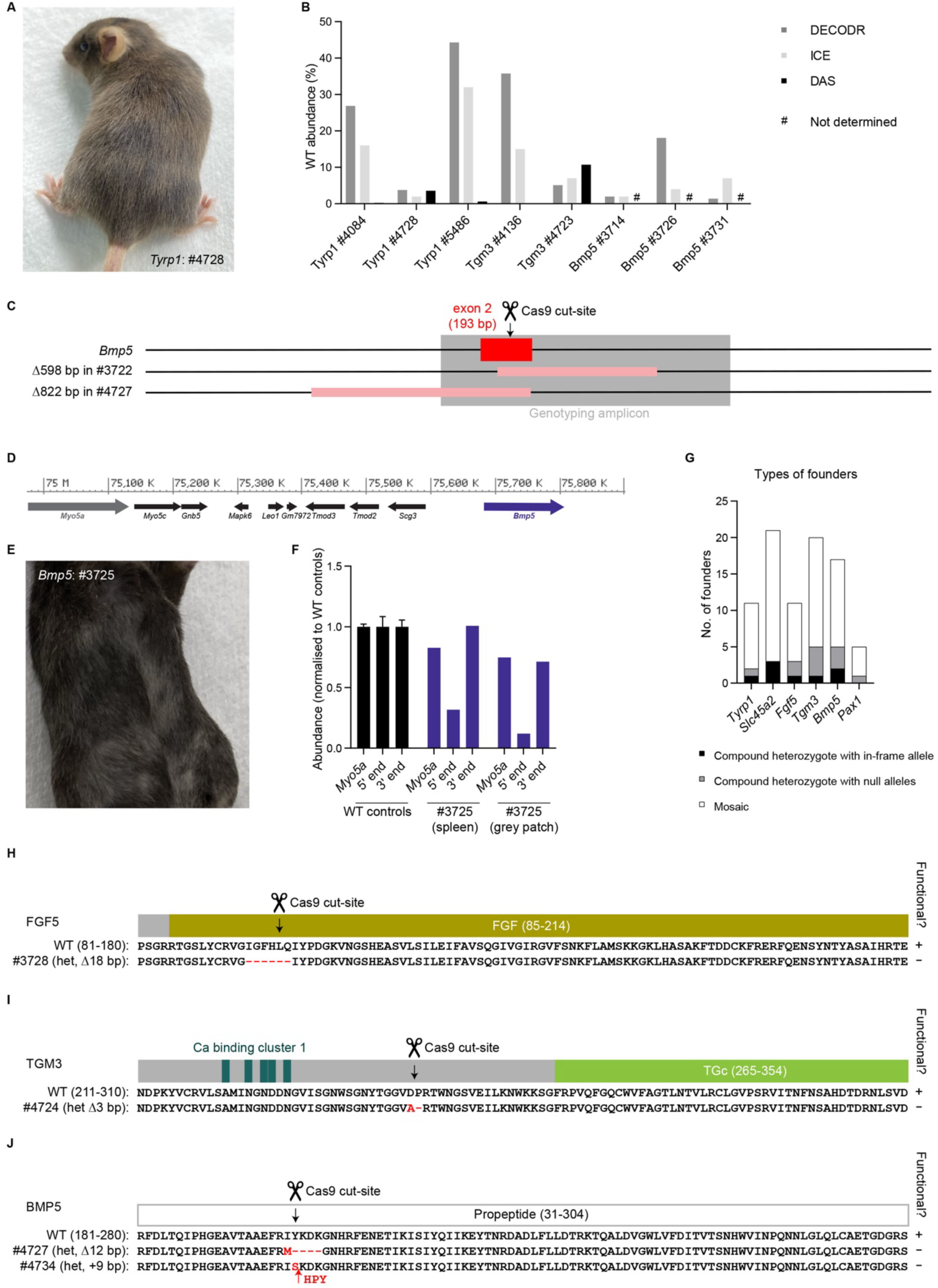
Compound heterozygous crispants enable genotype – phenotype associations. A. Photograph of *Tyrp1* founder #4728 showing the partial brown coat colour phenotype with black patches. B. Percentage of unedited sequence in all founders that had >0% unedited sequence predicted by Sanger deconvolution software (DECODR and ICE); and quantified by DAS in selected founders. C. Schematic of the large deletions in *Bmp5* founders, #3722 and #4727 (drawn to scale, deleted regions shown in pink). D. Schematic of the mouse chromosome 9 showing the *Myo5a* (“dilute”) gene 700 kb upstream of *Bmp5*. E. Photograph of *Bmp5* founder #3725 showing patches of grey fur. F. qPCR quantification at the upstream *Myo5a* locus and around the *Bmp5* 5’ and 3’ genotyping primer annealing sites in gDNA derived from the spleen and grey fur patch of founder #3725), normalised to three wild-type controls (mean ± SEM). G. Graph showing the number of founders generated for each gene that were compound heterozygous for an in-frame indel and frameshift/large deletion allele (black); compound heterozygous for two frameshift indels or one frameshift indel and one large deletion (grey); and with genetic mosaicism (white). H-J. Protein sequence schematic for the in-frame deletions in the compound heterozygous *Fgf5* (H), *Tgm3* (I) and *Bmp5* (J) founders. See Figure 3 for colour key.

### Crispants carry high levels of genetic editing and variable indel size in all genes

From our experiments detailed above, we generated a total of 60 founders with one targeted gene, 17 with one targeted endogenous gene plus EGFP, and six founders with two targeted endogenous genes. In total, we mutated the endogenous genes 89 times. Genotypic analysis of all five *Pax1* founders using DNA derived from tail (largely ectoderm), spleen (mesoderm) and liver (endoderm) showed high concordance between the three germ layers, with variation limited to the less abundant mutagenesis products (Figure S5; Table S3). This validated the use of tail tip DNA for genomic analysis in all other founders.

Phenotypic variability could result from incomplete editing. This is easily detectable in coat colour mutants (as black patches), but may not be observable in founders with the other visible phenotypes. Only one *Tyrp1* founder, #4728, had black (wild-type) patches (Figure 4A).

DECODR predicted 3.8% of its DNA was unmodified, which we confirmed by DAS (3.6%) (Figure 4B; Table S4). Since the phenotype is recessive, up to 7-8% of cells could retain TYRP1 protein function in this animal. DECODR predicted the presence of unedited sequence in one other *Tyrp1* founder (#5486). Like #4084 (discussed above), this animal displayed the full brown coat colour phenotype, and DAS revealed that it carried same 1 bp duplication that had been mis- assigned as unedited (Figure 1F, 4B, S1, S6A-D; Table S2, S5). DECODR predicted unedited sequence in two *Tgm3* founders (#4136 and #4723). DAS confirmed the presence of 10.7% unedited sequence in #4723 (Figure 4B, S6E-H; Table S4). In #4136, however, the deletion of one of a run of three cytosines (c.739delC; p.P247QfsX47) had been misassigned as unedited (Figure 4B, S6E-H; Table S4). Finally, a low level of unedited sequence was predicted in tail tip DNA in three *Bmp5* founders. As the short ear phenotype affects the mesoderm, we repeated this analysis using splenic DNA where no unedited sequence was detected in any of the founders (Figure 4B; Table S2). Overall, incomplete editing was only detected in two founders in this study.

We predominantly used DECODR to deconvolute Sanger sequencing chromatograms as it better predicted longer indels, but we found it grouped together two or more indels of the same length, assigning them as the most abundant sequence. In contrast, ICE separated these distinct repair products, as illustrated by *Pax1* founder #4088 (Figure S5D; Table S2). At all loci, we obtained a large number of short indels whose relative abundance reflected inDelphi predictions (Table S2-S5; Figure S5C). This was most clear in the *Tgm3* founders since this cut site had high microhomology strength, predicting an 8 bp deletion to occur with a frequency of 44.5%. This allele was detected in 10/20 *Tgm3* founders (Figure S6E-H, S7; Table S2-S5). Despite the preference for this repair product, no founders were homozygous for this *Tgm3* allele. We did not generate any homozygous founders (i.e. with the same mutation on both alleles).

Large on-target deletions were common. The longest deletion within the genotyping amplicons was Δ598 bp (554_683+368del in *Bmp5* founder #3722; Figure 4C; Table S2). ELF-CLAMP enabled us to sequence deletions that extended beyond the genotyping amplicons, the largest of which was Δ822 bp (c.485-634_672 in *Bmp5* founder #4727; Figure 4C; Table S4). The presence of grey patches on *Bmp5* (short ear) founder, #3725, indicated changes in the *Myo5a* gene, 700 kb away, in which recessive mutations cause a “dilute” coat colour (Figure 4D-E). We performed qPCR to determine whether deletions could extend to the *Myo5a* locus. There was a reduction at *Myo5a* in #3725 compared to wild-type controls, with lower levels in gDNA extracted from skin in the grey patches than from the spleen (Figure 4F). The only large insertion we detected was the 82 bp from intron 1 in *Slc45a2* founder #4062 (Figure 2G, S3A, mentioned above). Other types of previously reported genome damage (large inversions or mutations in cis with short indels) were not detected.

### Compound heterozygous crispant founders enable genotype – phenotype association

Several founders were compound heterozygotes (Figure 4G, S5, S7-9; Table S2-S5). Most of these (11-27% all founders per gene, except *Slc45a2*) carried two null alleles: two short frameshifting indels or one short frameshifting indel and one large deletion. For all genes except *Pax1*, we obtained at least one founder that was heterozygous for an in-frame indel (combined with a null allele). Since all traits are recessive, the phenotype of these founders reveal whether a short in-frame indel is sufficient to abolish protein function and therefore inform whether the mutated domain is required. As described above, these founders enabled us to determine that the tyrosinase domain of TYRP1 is critical for protein function and the first cytoplasmic domain of S45A2 is dispensable (Figure 1H, 2F; Table S2). The in-frame mutation in *Fgf5* founder #3728 (c.283_300del; p.I95_Q100del) demonstrated the functional importance of mutating the well-conserved FGF (fibroblast growth factor) domain (residues 85-214), involved in binding its receptor (Figure 4H, S9; Table S2). The in-frame mutation in *Tgm3* founder #4724 (c.737_739del; p.D246A,P247del) revealed that mutation of an uncharacterised region downstream of the first cluster of Ca^2+^ binding sites and upstream of the third transglutaminase domain impacts TGM3 protein function (Figure 4I, S9; Table S4). This mutation may specifically disrupt an unknown functional domain or it may impact TGM3 protein level, e.g. by disrupting splicing, post-translational processing or protein stability. The in-frame mutations in *Bmp5* founders #4727 (c.600_611del; p.I200M,Y201_K204del) and #4734: c.595_596 (insCTCATCCTT; p.Y199SinsHPY) revealed that mutation of the propeptide can affect BMP5 protein function, perhaps by affecting its cleavage (Figure 4J, S9; Table S4).

### Crispants can form an allelic series that shed light on protein domain function

We next asked if we could use this method to obtain non-null alleles that would shed light on protein domain function. For this, we aimed to produce truncated and in-frame alleles by targeting *Adamts20* (A disintegrin-like and metallopeptidase (reprolysin type) with thrombospondin type 1 motif, 20), associated with the “belted” phenotype. This gene is composed of 41 exons, alternatively spliced to form six isoforms. The most common isoform contains exons 3-41 (Figure 5A). ADAMTS20 protein consists of N-terminal Reprolysin and Disintegrin domains and 15 C-terminal thrombospondin type 1 (TSP) repeats. *Adamts20^bt^* is a missense mutation (c.1598C>T; p.L533P) in the Disintegrin domain^43^. Several other missense and nonsense mutations in *Adamts20* phenocopy *Adamts20^bt^*(Figure 5A). Of these, the best characterised truncated allele is *Adamts20^bt-Bei^*^1^ (c.2860C>T; p.R954X), which lies downstream of the disintegrin domain in the third TSP repeat^43^. We designed an sgRNA nearby R954X in exon 22 and produced 11 founders (Table 1, S1). Seven (63.6%) displayed the full white belt phenotype, three (27.3%) had a small patch of white fur and one (9.1%) appeared “wild-type” (Figure 5B, S4).

**Figure 5:**
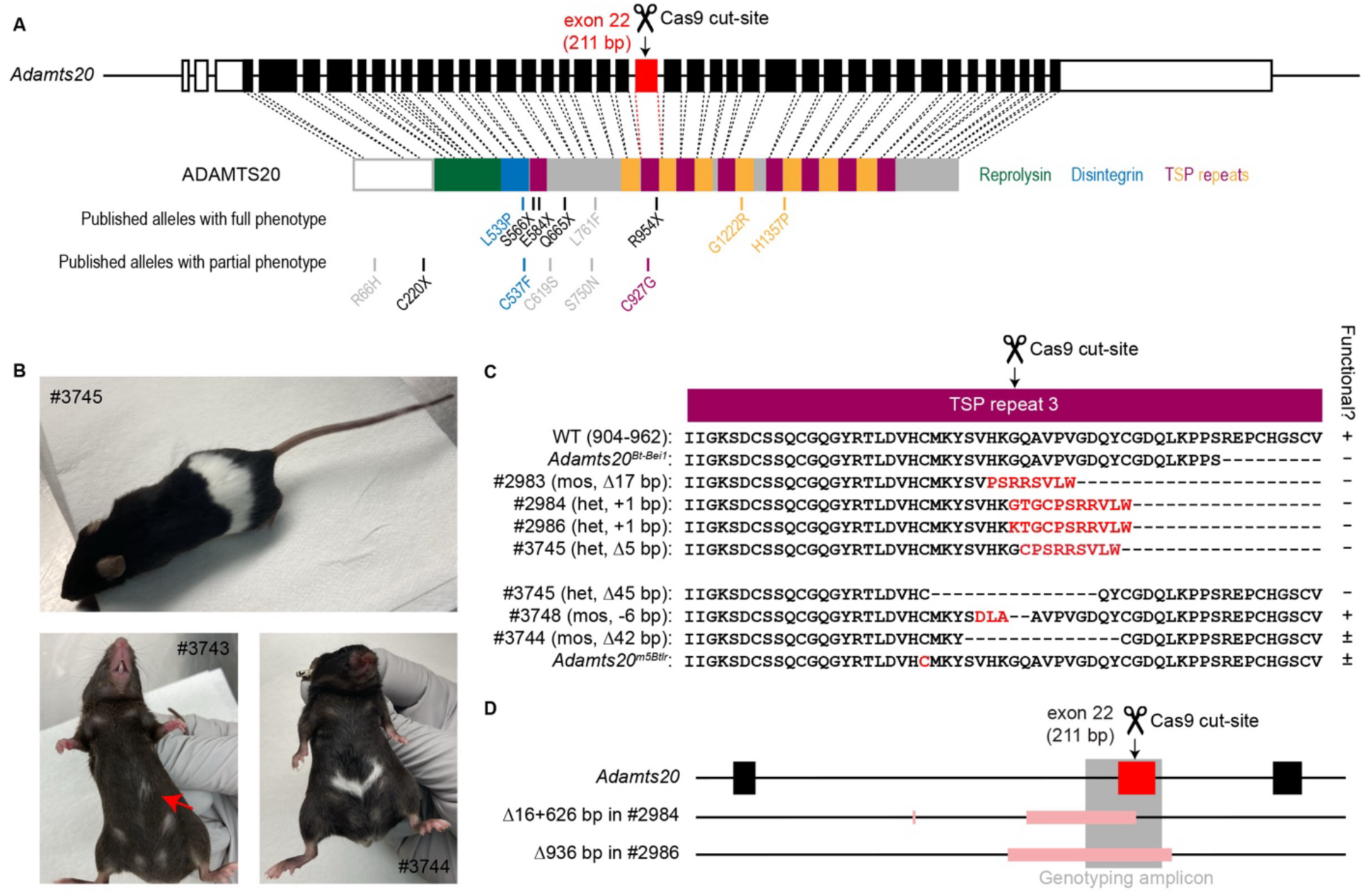
CRISPR/Cas9 mutagenesis can generate an allelic series of non-null mutations. A. Schematic of the *Adamts20* gene with the ADAMTS20 protein (exons and protein domains drawn to scale). The position of the Cas9 cut site is indicated by the scissors and arrow. Key: 5’ and 3’ UTRs: white; open reading frame (ORF): black; targeted exon: red; excised propeptide: white; Reprolysin: green; Disintegrin: blue; TSP repeats: purple and orange. Protein changes resulting from published alleles are shown below. Key: nonsense mutations: black; and missense mutations: colour matches domain. B. Photographs of selected founders showing the full belted phenotype (upper), partial belt from mosaicism (lower left) and partial belt from hypomorphic allele (lower right). C. Schematic of the protein sequence resulting from the truncating indels and in-frame deletions in compound heterozygous founders (“het”) and mosaic founders (“mos”) carrying one short indel and more than one large deletion; and the published *Adamts2^Bt-Bei1^*and *Adamts20^m5Btlr^* alleles. D. Schematic of the genome damage in the compound heterozygous founders, #2984 and #2986 (drawn to scale, deleted regions shown in pink).

Three of the seven founders with the full phenotype were compound heterozygotes. Two of these (#2984 and #2986) each carried a truncating indel and a large “unseen” deletion. Given that the trait is recessive, we can conclude that the truncated alleles are sufficient to cause the belted phenotype (Figure 5C, S4; Table S5), consistent with the *Adamts20^bt-Bei^*^1^ allele that we mimicked. We sequenced the “unseen” deletions by ELF-CLAMP. #2986 carried a 936 bp deletion, which was the largest deletion we observed in this study (Figure 5D; Table S6). #2984 carried a 626 bp deletion, with an additional 16 bp deletion 633 bp upstream (Figure 5D; Table S5). The third compound heterozygote (#3745) carried a long in-frame deletion (Δ45 bp): c.2782_2826del; p.M928_D942del and a frameshift mutation (Figure 5B-C, S10; Table S5), suggesting that mutating the TSP3 repeat is sufficient to abolish ADAMTS20 protein function.

The other four founders with the full phenotype were mosaic. One (#2983) carried a 17 bp deletion within the genotyping amplicon and several large deletions (sequenced using ELF-CLAMP). We predict that the large deletions are truncating/null since they span one/both ends of exon 22, so this exon will be excluded in *Adamts20* transcripts. This founder therefore carries one truncating allele and several null alleles (Figure 5C, S4; Table S5). The founder with the wild-type appearance (#3748) also carried one short mutation within the genotyping amplicon and several large deletions. This mutation was an in-frame (-6 bp) insertion-deletion (c.2795T>A, 2798_2807delinsTTGC; p.V932_336delinsDLA) (Figure 5C, S10; Table S6).

Unlike TYRP1 and S45A2 where loss-of-function phenotypes are cell-autonomous, ADAMTS20 protein is secreted^44^ and therefore it is not necessary for all cells to express functional ADAMTS20 for an animal to have a wild-type appearance. We therefore conclude that this short in-frame insertion-deletion in TSP3 does not disrupt ADAMTS20 protein function.

A partial phenotype could result from lower levels of secreted functional ADAMTS20 protein or from the presence of a hypomorphic variant. Two founders (#3734 and #3743) displayed small white patches, and carried multiple variant alleles including short in-frame indels (Figure 5B; Table S6). A third partially belted founder (#3744) carried a 42 bp deletion (c.2792_2833del; p.S931C, V932_C945del) and several large deletions (Figure 5B-C, S10; Table S6). We propose that the Δ42 bp allele is hypomorphic due to its size and the overlap with the non-functional 45 bp deletion in founder #3745 (Figure 5C). In support of this hypothesis, several published *Adamts20* alleles cause partial belted phenotypes, one of which lies in exon 22 and encodes a missense mutation in the TSP3 repeat (*Adamts20^m5Btlr^*: c.2779T>G; p.C927G; Figure 5A,C).

Using one sgRNA, we have produced eleven founders that include a non-null allelic series comprising three truncated alleles and three in-frame alleles with varying impact on coat colour that allowed us to classify them as loss-of-function, functional or hypomorphic.

## Discussion

Phenotypic assessment of crispant founders is replacing historical methods for genetic screening in zebrafish and *Xenopus* studies. Here, we demonstrate that crispant mouse founders display recessive traits and so could be adopted for initial *in vivo* assessment of mutant phenotypes. The major concern in the mouse community is that genetic mosaicism arising from mutagenesis occurring after cell division in zygotes will impact the phenotype^24,45,46^. Incomplete editing and the presence of short in-frame variants that retain function are of particular concern. By using recessive visible traits, we could determine whether individual founders displayed the expected phenotype. Incomplete editing was very rare: we achieved 100% editing in all but two animals in this study. Genetic mosaicism was common, but did not impact the phenotype when all alleles present disrupted function, including short in-frame indels.

The standard approach for generating null alleles is targeting a critical exon with an sgRNA^12^. Repair by NHEJ/MMEJ can generate frameshift mutations, which lead to the presence of premature termination codons (PTC) that target transcripts for nonsense mediated decay (NMD). This approach does not consider in-frame indels (which theoretically represent a third of repair products) to be sufficient to disrupt genes. Our results demonstrate that the best strategy to generate a null allele is to ensure that an sgRNA abolishes protein function by considering the protein domain structure and targeting an essential functional domain. We also show that targeting a domain of unknown function can reveal its functional importance in crispants. Short in-frame indels induced by the sgRNAs in this study disrupted protein function in *Tyrp1, Fgf, Tgm3* and *Bmp5*, but not in *Slc45a2* and *Adamts20*.

It is recommended to avoid targeting too close to the N-terminus of a gene because transcripts with a PTC <150 nt downstream of the start codon may also escape NMD, due to translation from a downstream in-frame start codon^47^. Our sgRNA targeting *Slc45a2* was within this range and there are three in-frame start codons just downstream. However, the presence of an ATG downstream of a PTC is not always sufficient to rescue gene function, as demonstrated by the null phenotype of the published mutation that we mimicked (*Slc45a2^uw-10Btlr^* c.39T>A; p.Y13X). *Slc45a2* founder #4062 revealed an independent mechanism by which targeting too close to the N-terminus could fail to disrupt a gene: mutagenesis can generate an alternative start codon, resulting in a variant where only the first few amino acids are lost.

On-target genome damage resulting from Cas9-mediated breaks is increasingly reported^32,33^. This includes large deletions, insertions and rearrangements. These events are often missed by routine genotyping assays: most commonly PCR amplification and sequencing of <1 kb regions around the target site. We detected the presence of large deletions by qPCR and sequenced a subset of these using ELF-CLAMP. When aiming to generate null alleles, genome damage is not an issue as long as it does not extend beyond the targeted gene. In *Bmp5* founder #3725, we detected deletions that extended to the *Myo5a* locus (700 kb away). We speculate, however, that extremely long deletions cannot occur at a high frequency as the loss of flanking genes will likely result in lethality - as is known for large deletions spanning *Myo5a* and *Bmp5*^48^.

Phenotypic assessment of crispants has several applications. While the phenotypes assessed here were detectable in individual animals, traits with lower penetrance could be assessed in cohorts of crispants (generated from a single round of mutagenesis). In situations when CRISPR/Cas9-mediated mutagenesis results in no live pups, it can be hard to determine whether this is due to technical failure or because the targeted gene is essential during embryogenesis^49^. The presence of developmental phenotypes in crispant embryos would suggest that the targeted gene is essential. Therefore, mouse lines could be generated by producing founders with incomplete editing (by reducing targeting efficiency or injecting CRISPR/Cas9 reagents into one blastomere in 2-cell embryos), which could be bred to generate heterozygotes. Similarly, when knocking out both copies of a gene causes infertility in both sexes, defects in gametogenesis could be detected in crispants, and mouse lines could instead be established using this strategy. Crispants can also be used for initial phenotypic assessment when investigating genetic interactions. IMPC mouse lines are generated on C57BL/6N and crispants could be produced on other strain backgrounds to assess for phenotypic modifiability. It is often desired to study the interaction between two genes. Unfortunately, we show that targeting both genes simultaneously dramatically reduces viability. Instead, we propose to establish/obtain a mouse line with a mutation in the first gene and use this line as donors to mutate the second gene in crispants. This strategy can be applied to screen for genetic suppressors of disease – which can reveal potential therapeutic targets^6–8^.

Making genotype-phenotype associations with non-null alleles helps to dissect protein function. We produced compound heterozygous crispants carrying an in-frame mutation and a null allele in five of the six genes that we targeted for disruption. For recessive traits, the presence or absence of the null phenotype in these founders revealed whether in-frame indels were sufficient to abolish protein function. C-terminal deletions can be used to assess the importance of the domain(s) downstream of a truncating mutation. As demonstrated here for *Adamts20*, truncating and in-frame variants can be generated using the same sgRNA. Compound heterozygous crispants could be bred to establish mouse lines with the alleles of interest for phenotypic confirmation and mechanistic studies. Mosaic animals carrying potentially informative variants could be also bred to segregate alleles for phenotypic analysis. We demonstrate that a single round of mutagenesis can generate many alleles and that these can be identified via thorough genotyping of crispant founders.

Using phenotypic assessment of crispants as a first stage in multiple experimental pipelines will inform on the most promising candidates to prioritise for further study. This will save considerable time and resources and dramatically reduce animal numbers.

## Materials and Methods

### Accession numbers for genes, transcripts and proteins

Nucleotide and amino acid numbering refer to the following accession numbers for mouse genes, transcripts and proteins. *Tyrp1* gene (NCBI ID: 22178,), *Tyrp1* transcript (NM_031202.3) and TYRP1 protein (P07147.1). *Slc45a2* gene (NCBI ID: 22293), *Slc45a2* transcript (NM_053077) and S45A2 protein (P58355.1). *Fgf5* gene (NCBI ID: 14176), *Fgf5* transcript (NM_010203.5) and FGF5 protein (P15656.1). *Tgm3* gene (NCBI ID: 21818), *Tgm3* transcript (NM_009374.3) and TGM3 protein (Q08189.2). *Bmp5* gene (NCBI ID: 12160), *Bmp5* transcript (NM_007555.4) and BMP5 protein (NP_031581.2). *Pax1* gene (NCBI ID: 18503), *Pax1* transcript (NM_008780.2) and PAX1 protein (NP_032806.2). *Adamts20* gene (NCBI ID: 223838), *Adamts20* transcript (NM_177431.5) and ADAMTS20 protein (P59511.2).

### Experimental model and subject details Mouse lines

Animal procedures were approved by the Animal Use Committee at the Canadian Council on Animal Care (CCAC)-accredited animal facility, The Centre for Phenogenomics (TCP). All mice were housed in a specific-pathogen-free (SPF) facility. They were maintained on a 12-h light/dark cycle and given ad libitum access to food and water. Mice were fed a standard diet (HarlanTeklad 2918), consisting of 18% protein, 6% fat and 44% carbohydrates. They were housed in individually ventilated cages (IVC) with wood chippings, tissue bedding and additional environmental enrichment in groups of up to five animals.

Wild-type C57BL/6J (The Jackson Laboratory) and Tg(CAG-EGFP)D4Nagy (JAX stock #018979; backcrossed onto C57BL/6J for >8 generations)^42^ were used to derive embryos. Wild-type or homozygous Tg(CAG-EGFP)D4Nagy females were used as embryo donors at 3-4 weeks. Wild-type or hemizygous Tg(CAG-EGFP)D4Nagy males were used for breeding to generate embryos. CD-1 (ICR) (Charles River Laboratories) outbred albino stock females were used as pseudopregnant recipients.

## Method details

### Generation of “crispant” founders

Crispant founder mice were generated by the Model Production Core at The Centre for Phenogenomics. Donor females were superovulated by intraperitoneal (IP) injection of 5 IU of pregnant mare serum gonadotropin (PMSG, Prospec, HOR-272) at 10 am (room light cycle: 5 am/on, 5 pm/off) followed 48 hours later by an IP injection of 5 IU human chorionic gonadotrophin (hCG, EMD Millipore 230734) and mated overnight with proven breeder males. The next morning the females were checked for the presence of a vaginal copulation plug as evidence of successful mating. Oviducts were dissected at approximately 22 hours post hCG and cumulus oocyte complexes were released in M2 medium (Millipore MR-015D) and treated with 0.3 mg/ml hyaluronidase (Sigma H4272) as previously described^50^. Fertilized embryos were selected and kept at 37°C, 6% CO_2_ in microdrops of KSOM media with amino acids (KSOM^AA^, Life Global medium LGGG-050 from Cooper Surgical) covered by embryo tested paraffin oil (Life Global LGPO-500 from Cooper Surgical) prior to electroporation. Embryos were briefly cultured in KSOM^AA^ and transferred into the oviducts of 0.5 dpc pseudopregnant CD-1(ICR) female recipients shortly after manipulations^50^.

### Reagents for Cas9 RNA-guided nuclease mediated mutations

To produce null alleles, single guide RNAs (sgRNAs) were designed to target exons that had previously been deleted or mutated with a truncating mutation in knock-out mouse lines (Table S1). The *Adamts20* sgRNA was designed to target nearby the truncating mutation, *Adamts20^bt-Bei1^* (c.2860C>T; p.R954X)^43^. sgRNAs were designed using CRISPOR (http://crispor.tefor.net/)^51^ and CHOPCHOP (https://chopchop.cbu.uib.no/)^52^ with off-target PAM set to -NRG. Off target criteria: at least three mismatches to unintended targets, at least one of which was in the 11 bp “seed” region adjacent to the PAM. All sgRNA had a predicted frameshift frequency >67%, assessed using inDelphi^26^. See Table S7 for sgRNA sequences. Recombinant sgRNAs (Synthego) and Cas9 protein (IDT) were used for electroporation. Mixes consisted of Cas9 protein (6 µM) and sgRNA (9 µM total or 4.5 µM each for 2 sgRNAs) in 1X buffer (100 mM KCl, 20 mM Hepes, pH 7.2-7.4) prior to dilution in an equal volume of OptiMEM (ThermoFisher 31985062) immediately before electroporation.

### Electroporation

Electroporation was performed as described^12,14^. Briefly, zygotes were treated with Acid Tyrode’s (Sigma T1788) for a few seconds to weaken the zona pellucida, rinsed through Opti-MEM media (Thermo Fisher Scientific 31985062) and placed into 5 µl 1:1 mixture of Cas9 RNP and Opti-MEM between electrodes on 1-mm gap electrode slide BEX CUY21 EDIT II connected to BioRad Gene Pulser XCell electroporator. Twelve square pulses at 30 V with 1 ms pulse duration and 100 ms interval were applied. The zygotes were retrieved from the slide, rinsed with KSOM^AA^ and kept at 37°C, 6% CO2 until ready for embryo transfers.

### Genomic DNA extraction

For initial PCRs, DNA was extracted from tail tips by incubation in alkaline lysis buffer (25 mM NaOH, 0.2 mM EDTA) at 95°C for 30 minutes, followed by neutralisation with an equal volume of 40 mM Tris-HCl. For additional assays, DNA was purified from ∼1 cm tail after termination by digestion in 500 μl lysis buffer (50 mM Tris HCl pH 8.0, 100 mM NaCl, 50 mM EDTA, 0.5% SDS) with 0.2 mg/ml Proteinase K overnight at 55°C. Samples were then treated with 0.1 mg/ml RNase at 37°C for 1-2 hours and gDNA was extracted with 500 μl PCI (phenol:chloroform:isoamyl alcohol, Sigma). gDNA was precipitated from the aqueous phase with 1 volume cold isopropanol, and dissolved in TE. DNA was purified from snap-frozen half spleen and portion of liver tissue by homogenisation in 3 ml lysis buffer (50 mM Tris HCl pH 7.5, 100 mM NaCl, 5 mM EDTA). Proteinase K was added to a final concentration of 0.4 mg/ml and SDS to 1% (w/v). After overnight incubation at 55°C, samples were treated with 0.1 mg/ml RNase at 37°C for 1-2 hours and gDNA was extracted with 3 ml PCI (phenol:chloroform:isoamyl alcohol, Sigma). gDNA was precipitated from the aqueous phase with 2.5 volumes 100% ethanol and 0.1 volumes 3M NaOH, and dissolved in TE.

### Genotyping PCRs

Tail tip genomic DNA was PCR amplified for genotyping. From *Pax1* founders, gDNA was also derived from spleen and liver. From selected *Bmp5* founders, gDNA was also derived from spleen. Quanta polymerase was used for all reactions and amplicon lengths were measured on a QIAxcel Advanced Instrument. Amplicons were Sanger sequenced using a forward nested primer for *Tyrp1*, *Slc45a2*, *Fgf5*, *Tgm3*, *Pax1* and *Adamts20* and a reverse nested primer for *Bmp5* (a polyT run upstream of the target site prevents the use of a forward primer for Sanger sequencing). See Table S8 for primer sequences and annealing temperatures.

### Sanger deconvolution analysis

Sanger deconvolution was performed using DECODR v3.0^29^ and ICE (Synthego)^30^. Long in-frame indels were defined as >21 bp (consistent with ICE). Sequence alignments between Sanger chromatograms and the wild-type reference were performed using Snapgene (Version 6.1.2).

### qPCR

Quantification of DNA copy number around genotyping primer annealing sites was performed by qPCR using gDNA diluted to 2.5 ng/µl. Each region of interest (ROI) was detected by region-specific primers and normalised to the TRFC locus (see Table S8 for primer sequences). qPCR was performed on a Viia7 instrument (ABI) instrument using SYBR Green PCR Master Mix (Invitrogen). PCR amplification conditions: annealing temperature 58°C, 40 cycles. Relative abundance = E^TFRC^^CT^TFRC^ / E^ROI^^CT^ROI^. Abundance was normalised to the mean of three wild-type animals.

### Deep-amplicon sequencing (DAS)

Genotyping PCR amplicons were sent for Premium PCR sequencing at Plasmidsaurus. The sequencing yield was 2914 – 4201 reads per sample. FASTQ format data were analysed using the CRISPResso2 software^53^ with the following parameters: Cas9, single end reads, 50% minimum homology. 1501 - 2672 reads per sample aligned and were included in the analysis. For quantification, the editing window was 3 bp either side of the Cas9 cut site. Indels observed in >0.5% of reads were plotted.

### Enrichment of Long DNA Fragments using Capture and Linear Amplification (ELF-CLAMP)

This protocol was adapted from Chang et al.^31^ for use with genomic DNA (gDNA). This protocol consists of two selection steps for viewpoints of interest (*in-vitro* CRISPR/Cas9 cutting and site-specific fusion of a biotinylated T7 promoter) followed by specific enrichment using linear amplification (*in-vitro* transcription). The resulting RNA is subsequently analysed using direct-RNA sequencing on an Oxford Nanopore Technologies MinION device. The samples were run in three batches.

sgRNAs for *in vitro* CRISPR/Cas9 cutting of viewpoints in the genomic DNA were produced by *in vitro* transcription of a dsDNA generated by annealing a ssDNA containing a T7 promoter (TAATACGACTCACTATAGGGAG), the specific sequence of the gRNA and a part of the common gRNA sequence (GTTTTAGAGCTAGAAATA), to a reverse oligo containing only the common sequence (see Table S9 for oligo sequences). This partially annealed template is next made fully double stranded using DNA Polymerase I, Large (Klenow) Fragment (New England Biolab, M0210). *In vitro* transcription of the resulting template to generate full-length gRNAs was performed by using the T7 RiboMAX Express Large Scale RNA Production System (Promega, P1320) followed by DNase treatment, according to the manufacturer’s instructions. sgRNAs were then isolated by 1:1 phenol-chloroform-IAA extraction followed by isopropanol with sodium acetate precipitation.

CRISPR/Cas9 cutting of viewpoints in genomic DNA was done as follows: 999 ng each gRNA was incubated with 30.5 pmol Alt-R S.p. Cas9 Nuclease V3 (Integrated DNA Technologies) in a total of 2.5 μl NEBuffer 3.1 (New England Biolabs) at room temperature for 10 minutes. 2.5 μg genomic DNA was combined with the sgRNA-Cas9 complexes (RNPs) in a total volume of 25 μl (in nuclease-free water) and incubated overnight at 37°C, followed by enzyme deactivation at 65°C for 30 minutes. Nuclear RNA and gRNAs were removed by adding 125 U RNase If (New England Biolabs, M0243) and incubation at 37°C for 45 minutes, followed by enzyme deactivation at 70°C for 20 minutes. The cut genomic DNA was purified and concentrated by adding 1 volume AMPure XP beads (30 μl) (Beckman Coulter, A63880) and eluted in 23 μl nuclease-free water. To repair single stranded damage, 20 μl of cut genomic DNA was mixed with 2.5 μl NEBNext FFPE Repair Buffer and 0.5 μl NEBNext FFPE Repair Mix (New England Biolabs, M6630), followed by incubation at 20°C for 15 minutes and addition of 3 volumes of AMPure XP beads (70 μl) for purification. DNA was eluted in 30 μl nuclease-free water.

Site-specific fusion of a biotinylated T7 promoter was done by adding probes on one side or both sides of the newly generated cut site. Probes consisted of the following components: a first biotinylated base, a short linker sequence, the recognition site for the *Sbf*I restriction enzyme (CCTGCA^GG), the complete T7 promoter sequence (TAATACGACTCACTATAGGGAG) and a 30 bp sequence that is complementary to the sequence directly bordering the CRISPR/Cas9 cut site (see Table S9 for probe sequences). Biotinylated probes were diluted to 1 μM by mixing 2 μl of probes with nuclease-free water in a final volume of 80 μl. This was added to 28.5 μl of cut and repaired genomic DNA, 10 μl of 5X OneTaq buffer, 1 μl of 10 mM dNTPs and 1 μl of OneTaq Polymerase (New England BioLabs, M0480). To generate (partially) double stranded DNA, the reaction was incubated in a thermal cycler with the following steps: 95°C for 8 minutes, 1°C decrease per 15 sec to 65°C, 68°C for 5 minutes, rapid decrease to 4°C. Nuclease-free water was subsequently added to a final volume of 200 μl. Samples were pooled here for downstream processing: in each experimental batch, all samples with proximal probes were pooled and all samples with distal probes were pooled.

5 μl per probe of Dynabeads MyOne Streptavidin C1 beads (Thermo Fisher Scientific, 65001) were washed according to the manufacturer’s instructions, followed by resuspension in 200 μl of Binding&Wash buffer (10 mM Tris-HCl, pH 7.0, 1mM EDTA, 2M NaCl). The total volume of beads was added to the cut and T7 promoter-fused genomic DNA, followed by incubation at room temperature for 30 minutes in a HulaMixer (Thermo Fisher Scientific). Beads were washed three times in 200 μl of 1X Binding&Wash buffer according to the manufacturer’s instructions, and further washed in 43 μl nuclease-free water with 5 μl CutSmart Buffer at 37°C for 20 minutes. Bound DNA fragments were released by adding 43 μl nuclease-free water, 5 μl CutSmart Buffer, and 20 U *Sbf*I (New England BioLabs, R3642), followed by incubation at 37°C for 20 minutes. The released DNA was purified and concentrated by adding 1 volume AMPure XP beads and eluted in 11 μl nuclease-free water.

10 μl of the eluted T7 promoter-fused DNA was *in vitro* transcribed using the T7 RiboMAX Express Large Scale RNA Production System (Promega, P1320) according to the manufacturer’s instructions with minor modifications: 10 μl of libraries were added to 15 μl 2x T7 RiboMax buffer, 3 μl T7 Enzyme Mix, 2 μl 5M Betaine (Sigma-Aldrich, B0300) and 1 μl SUPERase RNase Inhibitor (Thermo Fisher Scientific, AM2694) and incubated at 37°C for 60 minutes. In each batch, the proximal and distal samples were now combined. The RNA was poly(A) tailed using Poly(A) Tailing Kit (Thermo Fisher Scientific, AM1350) according to the manufacturer’s instructions and incubated at 37°C for 10 minutes. The poly(A) tailed RNA was purified and concentrated by adding 100 μl of Agencourt RNAClean XP beads (Beckman Coulter, A63987) and eluted in 12 μl nuclease-free water. RNA concentration was determined using the RNA HS Assay on a Qubit Fluorometer (Thermo Fisher Scientific, Q32852).

Nanopore direct-RNA sequencing libraries were prepared using the Direct-RNA Sequencing Kit, version SQK-RNA002 (Oxford Nanopore Technologies) according to the manufacturer’s instructions with minor modifications: both adaptor ligation steps were performed for 15 minutes and the reverse-transcription step was performed at 50°C for 30 minutes with 1 μl of SUPERase RNase Inhibitor supplemented. The final Agencourt RNAClean XP beads purification step was done using 24 μl of beads for a stringent size selection while maintaining enough yield.

Direct-RNA sequencing was done for 72 hours, using FLO-MIN106 R9.4.1 flowcells on a MinION (MK 2.0) sequencing device with the MinKNOW software (Oxford Nanopore Technologies).

Direct-RNA sequencing reads (POD5 files, converted from FAST5) were basecalled with Dorado v0.6.1 using the high-accuracy (“hac”) model without read trimming. FASTQ reads were binned by viewpoint using Cutadapt, retaining only those with the expected viewpoint sequence at the 5’ end. A subsequent filtering step excluded off-target ELF-CLAMP products by requiring the presence of the corresponding downstream gene sequence. Filtered reads were then aligned to the soft-masked *Mus musculus* reference genome (GRCm39) using BWA-MEM. For visualisation, SAM files were converted to BAM format and loaded into the Integrative Genomics Viewer (IGV). All analysis scripts are available at https://github.com/fbm48/Tillotson, and the basecalled FASTQ files are deposited in the NCBI GEO under accession number GSE289584.

Alignments using SAM files were analysed to determine the sequence of large “unseen” deletions, and to check for the presence of genome damage in regions flanking short in-frame indels (in compound heterozygous founders). Deletions detected by ELF-CLAMP were confirmed by PCR amplification and Sanger sequencing (see Table S8 for primer sequences and annealing temperatures).

## Statistical analysis

Percentage viability was compared by Fisher’s exact tests using GraphPad Prism 9. Significance thresholds: not significant (NS) P > 0.05; * P < 0.05; ** P < 0.01; *** P < 0.001.

## Acknowledgements

This work was funded by a Sir Henry Wellcome Postdoctoral Fellowship (210913/Z/18/Z) to R.T. and a Canadian Institutes of Health Research Foundation Grant (FDN-15423) to M.J.J.. R.T. is currently a University of Edinburgh Chancellor’s Fellow. L.-H. C. was funded by the Blood and Transplant Research Unit in Precision Cellular Therapeutics. The authors wish to acknowledge Sandra Tondat, Amit Patel and Maribelle Cruz in the TCP Model Production Core for their contribution to the generation of the crispant founders, and members of the Justice laboratory, James Davies, Grzegorz Kudla, and Duncan Sproul for helpful discussions.

## Author contributions

Conceptualisation, R.T., M.J.J. and L.M.J.N.; methodology, R.T., M.G., L-H.C. and L.G.L.; formal analysis, R.T., F.B.M. and P.G.; investigation, R.T., J.R. L-H.C. and C.T.; writing – original draft, R.T.; writing –review & editing, all of the authors; visualisation, R.T. and J.R.; supervision, M.J.J., and L.M.J.N.; project management, J.R.; funding acquisition, R.T. and M.J.J.

---

**Figure S1:**
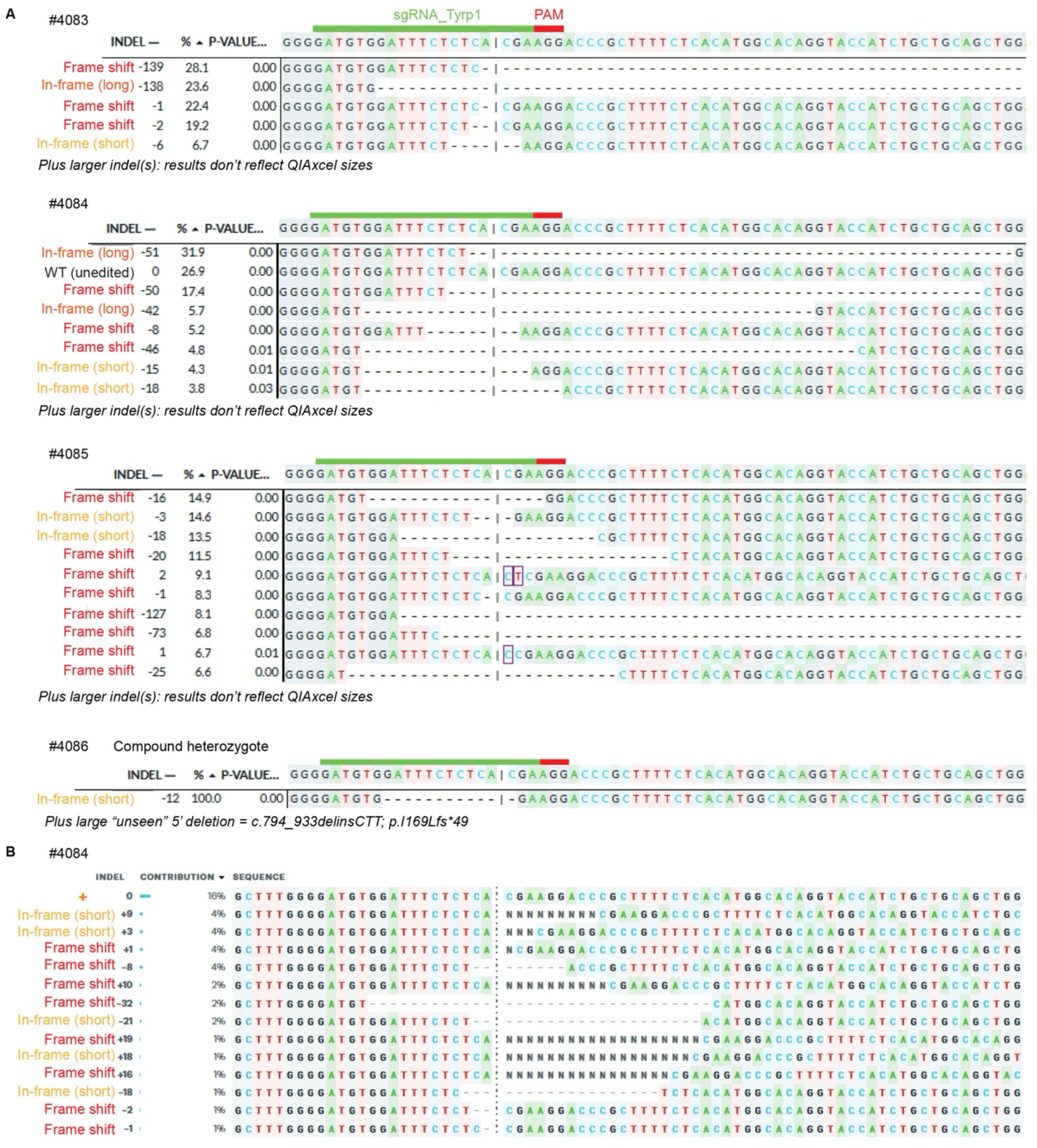
Genotyping analysis of *Tyrp1* founders. A. DECODR deconvolution of Sanger sequencing chromatograms of the genotyping amplicon in *Tyrp1* founders. B. ICE deconvolution of Sanger sequencing chromatogram of the genotyping amplicon in *Tyrp1* founder #4084.

**Figure S2:**
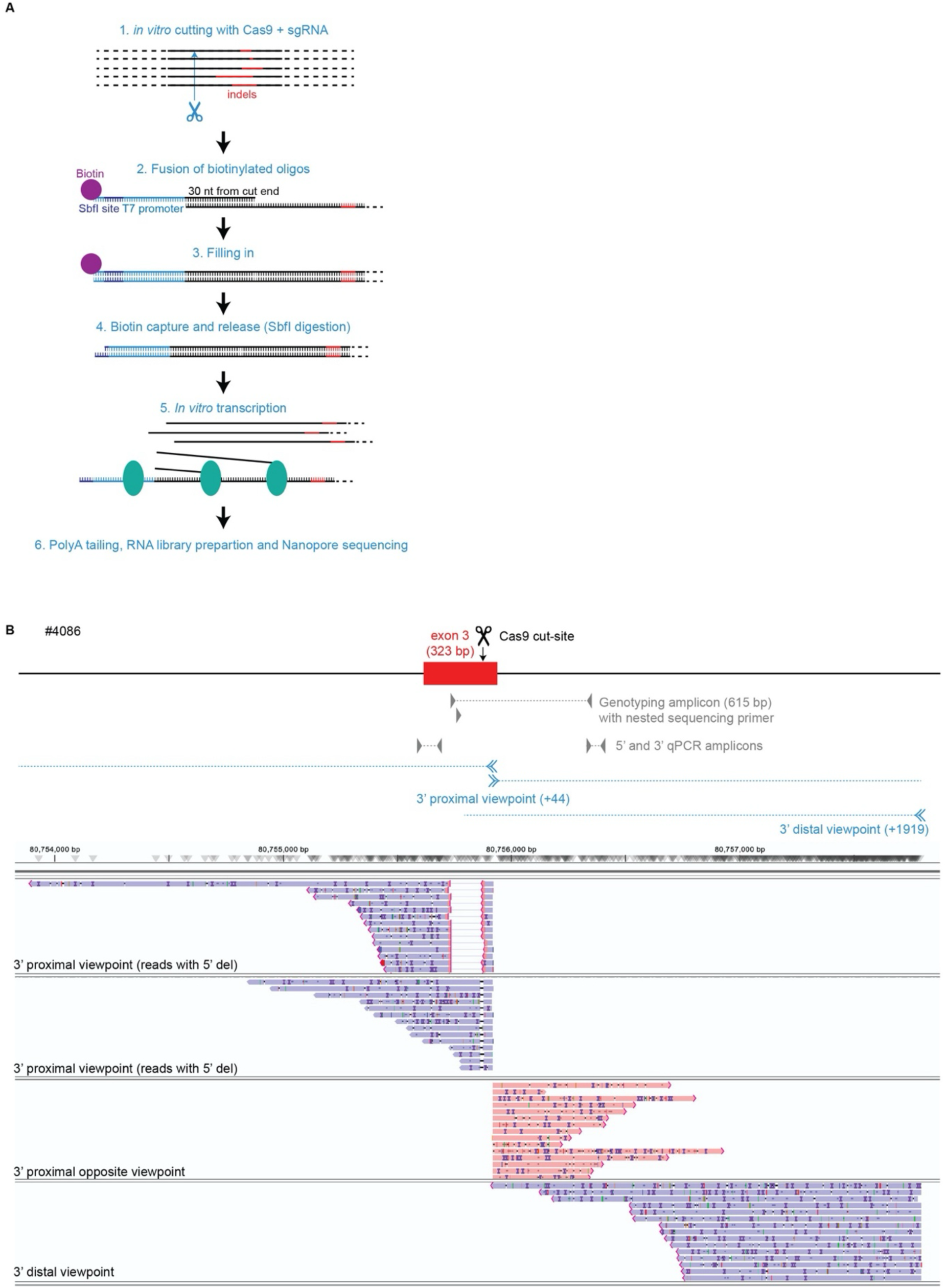
ELF-CLAMP analysis of the compound heterozygous *Tyrp1* founder. A. Schematic of the ELF-CLAMP experimental workflow. B. Screenshot of ELF-CLAMP sequencing reads viewed in IGV for *Typr1* founder #4086 are shown below a schematic of the *Tyrp1* gene showing the location of the Cas9 cut site, genotyping and qPCR primers and the viewpoints used for ELF-CLAMP.

**Figure S3:**
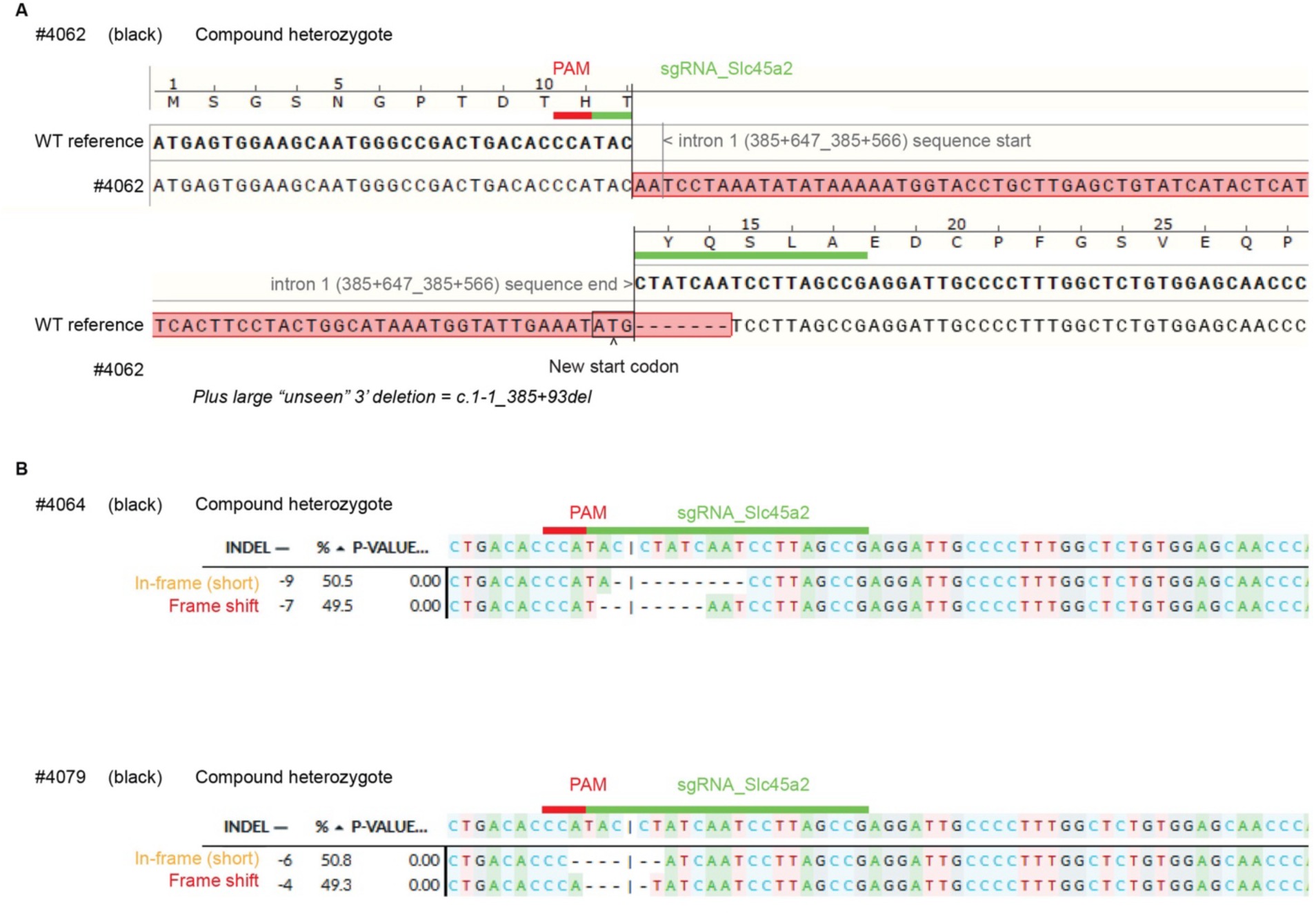
Genotyping analysis of *Slc45a2* founders. A. Sequence alignment of the complex deletion-insertion present in *Slc45a2* founder #4062 compared to the wild-type (reference) sequence. B. DECODR deconvolution of Sanger sequence chromatograms of the genotyping amplicon in *Slc45a2* founders #4064 and #4079.

**Figure S4:**
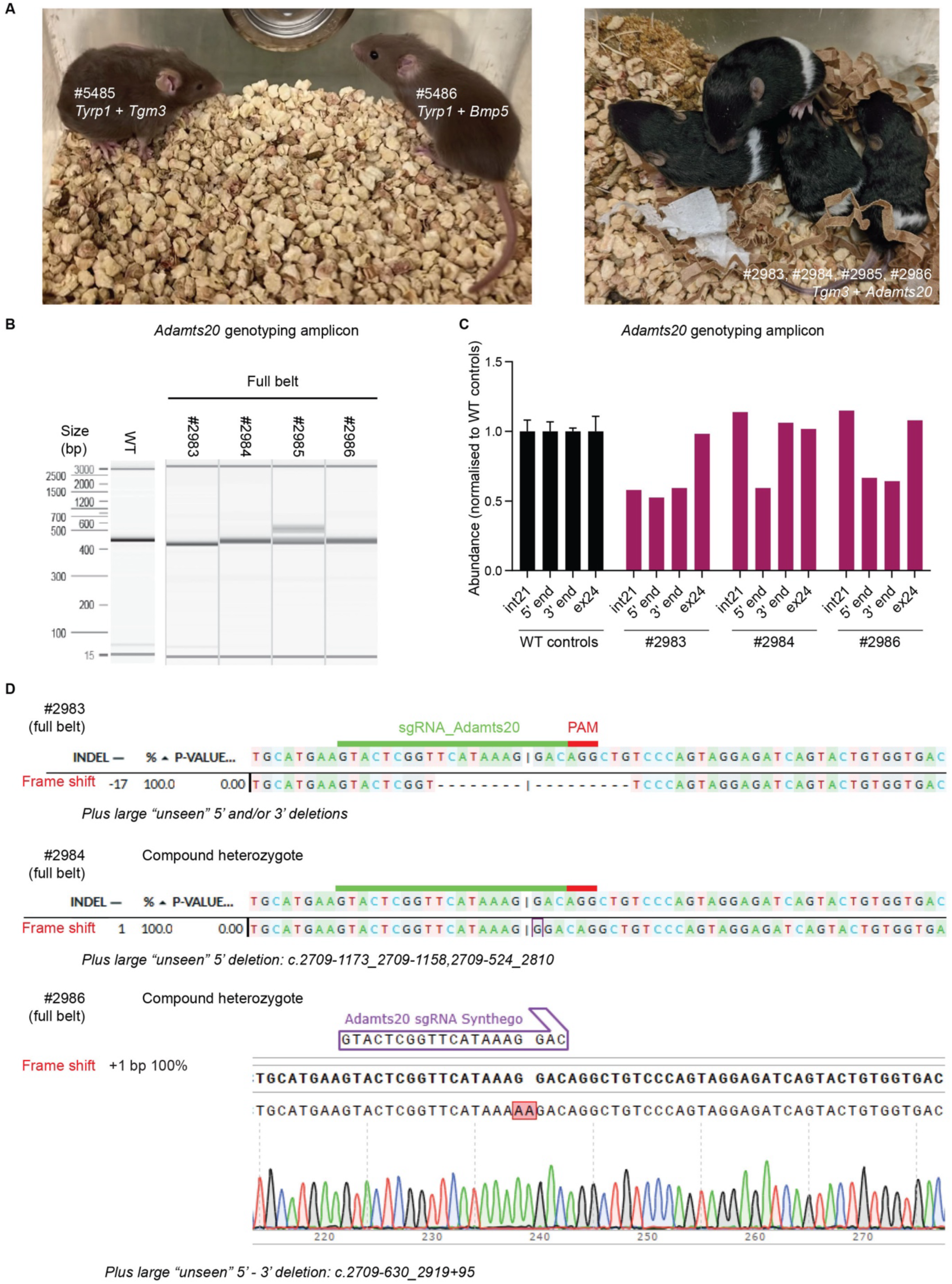
Crispant mice with two mutated genes display both phenotypes. A. Photographs of founders in which two endogenous genes have been mutated. From left: #5485 (*Tyrp1* + *Tgm3*), #5486 (*Tyrp1* + *Bmp5*) and #2983-6 (*Tgm3* + *Adamts20*). B. PCR amplification around the *Adamts20* target site in all four *Tgm3* + *Adamts20* founders, beside a wild-type reference (437 bp). C. qPCR quantification around the 5’ and 3’ *Adamts20* genotyping primer annealing sites and more distal sites in intron 21 and exon 24 (∼1.5 kb away from the targeted site) in founders #2983, #2984 and #2986), normalised to three wild-type controls (mean ± SEM). D. DECODR deconvolution of Sanger sequence chromatograms of the *Adamts20* genotyping amplicon in founder #2983 (mosaic for c.2798_2814del; p.H933PfsX9 and large 5’ and/or 3’ deletions) and #2984 (compound heterozygous for c.2804_2805insG; p.G935GfsX13 and a large 5’ deletion) and sequence alignment of the *Adamts20* genotyping amplicon in founder #2986 (compound heterozygous for c.2803delGinsAA; p.G935KfsX13 and a large 5’ - 3’ deletion).

**Figure S5:**
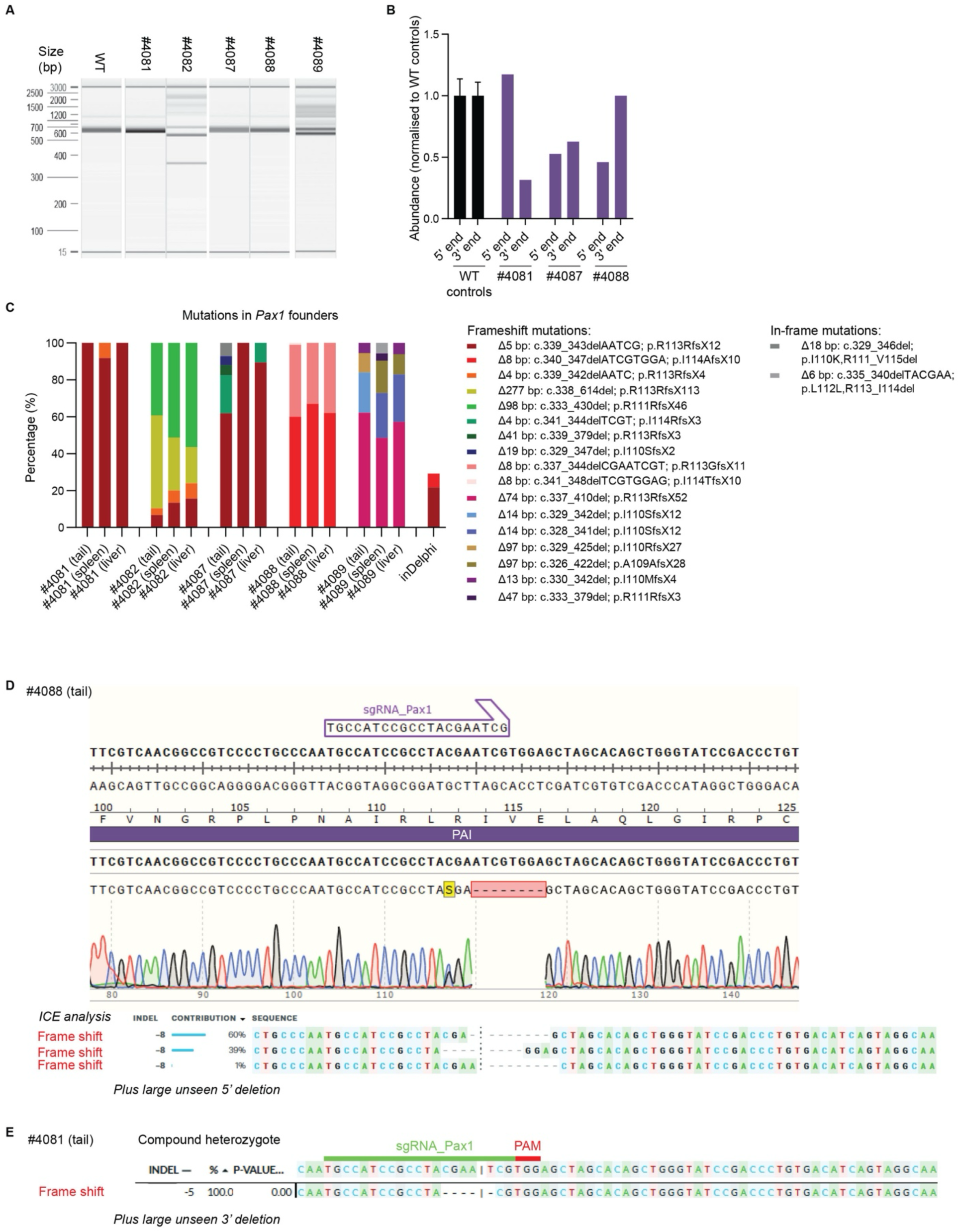
Mutations show high concordance across germ layers. A. PCR amplification around the targeted site in all five *Pax1* founders (tail), beside a wild-type reference (630 bp). B. qPCR quantification around the 5’ and 3’ genotyping primer annealing sites in founders #4081, #4087 and #4088), normalised to three wild-type controls (mean ± SEM). C. Genotypic analysis of all five *Pax1* founders using DNA derived from tail (largely ectoderm), spleen (mesoderm) and liver (endoderm). Graph shows the percentage of each repair product. inDelphi predicted two products, plotted by predicted frequency (percentage). D. Sequence alignment and ICE deconvolution of Sanger sequence chromatogram from the genotyping amplicon in founder #4088, using tail gDNA. E. DECODR deconvolution of Sanger sequence chromatogram of founder #4081 (compound heterozygous for c.339_343del; p.R113RfsX12 and a large 3’ deletion), using tail gDNA.

**Figure S6:**
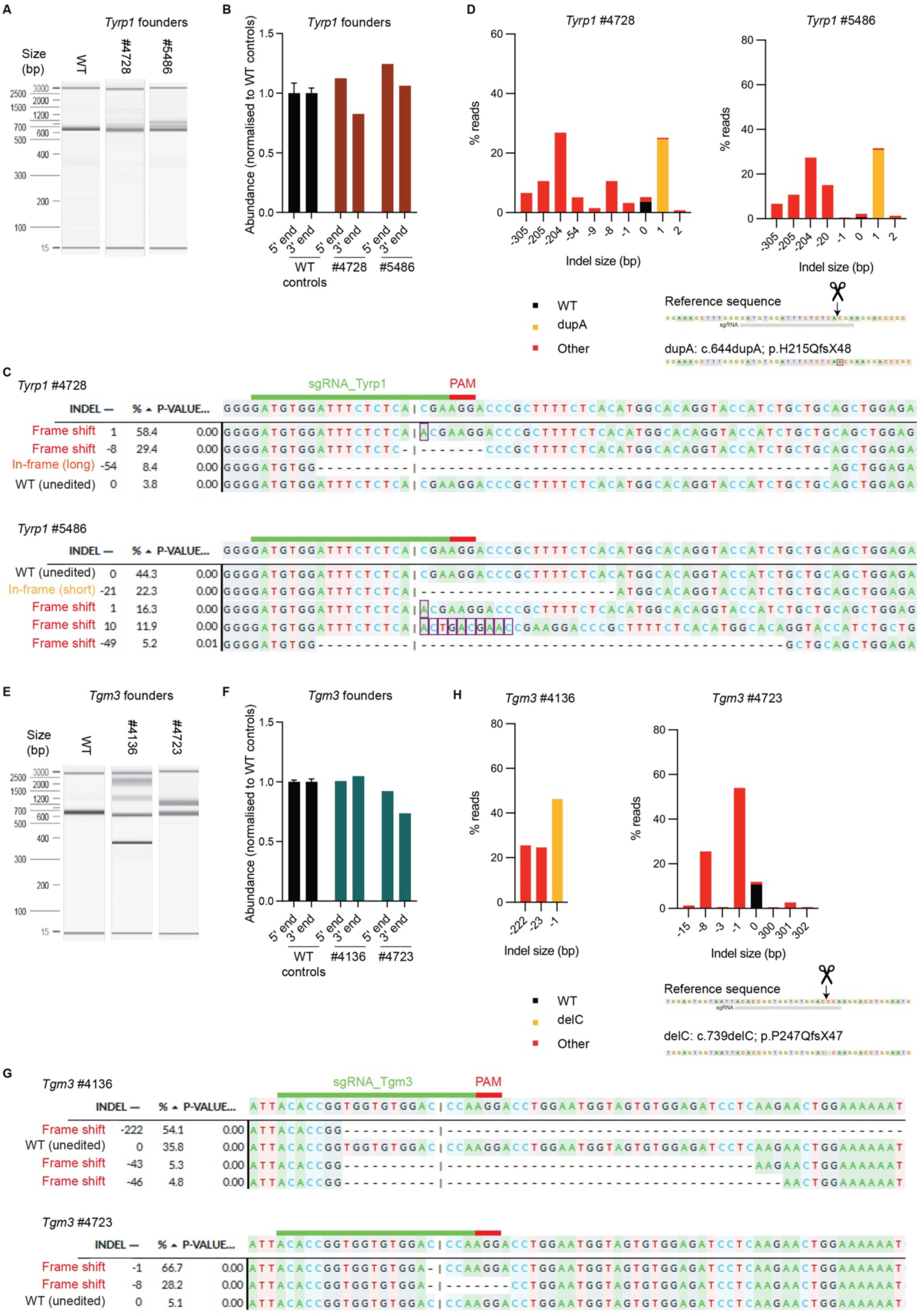
Genotyping analysis of *Tyrp1* and *Tgm3* founders with predicted unedited sequence. A. PCR amplification around the targeted site in the two additional *Typr1* founders with predicted unedited sequence (#4728 and #5486), beside a wild-type reference (615 bp). B. qPCR quantification around the 5’ and 3’ genotyping primer annealing sites in founders #4728 and #5486, normalised to three wild-type controls (mean ± SEM). C. DECODR deconvolution of Sanger sequence chromatograms of the genotyping amplicon in the *Tyrp1* founders. D. DAS analysis of #4728 and #5486 (percentage abundance of indels present in >0.5% reads are shown). E. PCR amplification around the targeted site in the two *Tgm3* founders with predicted unedited sequence (#4136 and #4723), beside a wild-type reference (609 bp). F. qPCR quantification around the 5’ and 3’ genotyping primer annealing sites in founders #4136 and #4723, normalised to three wild-type controls (mean ± SEM). G. DECODR deconvolution of Sanger sequence chromatograms of the genotyping amplicon in *Tgm3* founders. H. DAS analysis of #4136 and #4723 (percentage abundance of indels present in >0.5% reads are shown).

**Figure S7:**
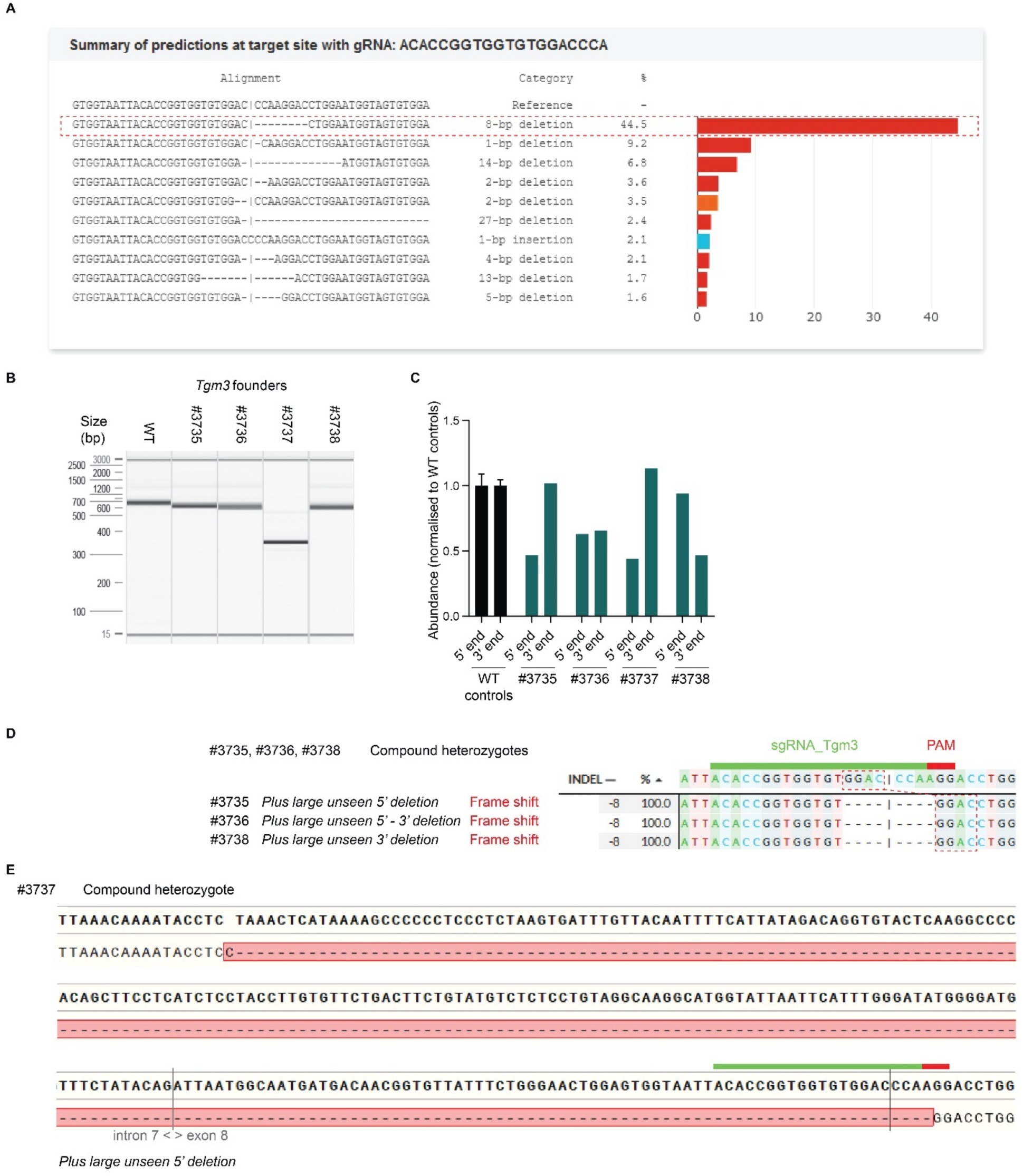
High microhomology of *Tgm3* cut site results in founders that are heterozygous for the predicted repair product. A. InDelphi prediction of repair products at the *Tgm3* cut site. High microhomology promotes an 8 bp deletion: c.739-746del p.P247LfsX4. B. PCR amplification around the targeted site in the four compound heterozygous *Tgm3* founders (#3735, #3736, #3737 and #3738), beside a wild-type reference (609 bp). C. qPCR quantification around the 5’ and 3’ genotyping primer annealing sites in these founders, normalised to three wild-type controls (mean ± SEM). D. DECODR deconvolution of Sanger sequence chromatograms of the genotyping amplicon in founders #3735, #3736 and #3738. E. Sequence alignment of Sanger sequencing of the genotyping amplicon in founder #3737.

**Figure S8:**
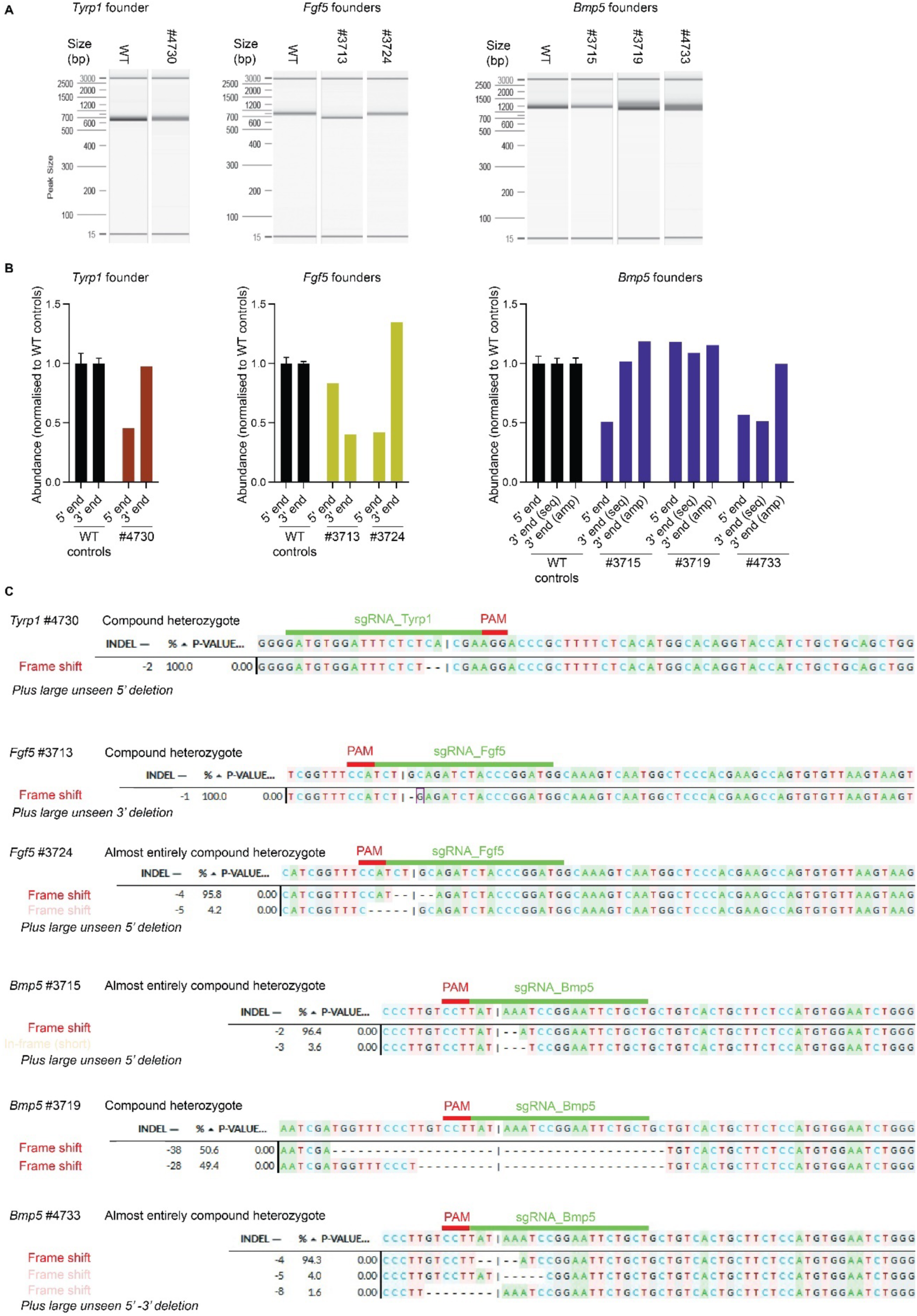
Genotyping analysis of compound heterozygous *Tyrp1*, *Fgf5* and *Bmp5* founders carrying two knock-out alleles. A. PCR amplification around the targeted site in compound heterozygous founders carrying two knockout alleles: *Tyrp1* #4730; *Fgf5* #3713 and #3724; and *Bmp5* #3715, #3719 and #4733. B. qPCR quantification around the 5’ and 3’ genotyping primer annealing sites (and the nested sequencing primer annealing site for *Bmp5*) in founders, normalised to three wild-type controls (mean ± SEM). C. DECODR deconvolution of Sanger sequence chromatograms of the genotyping amplicon in these founders. For genotyping analysis of *Pax1* founder #4081, see Figure S5; and of *Tgm3* founders #3735, #3736, #3737 and #3738, see Figure S7.

**Figure S9:**
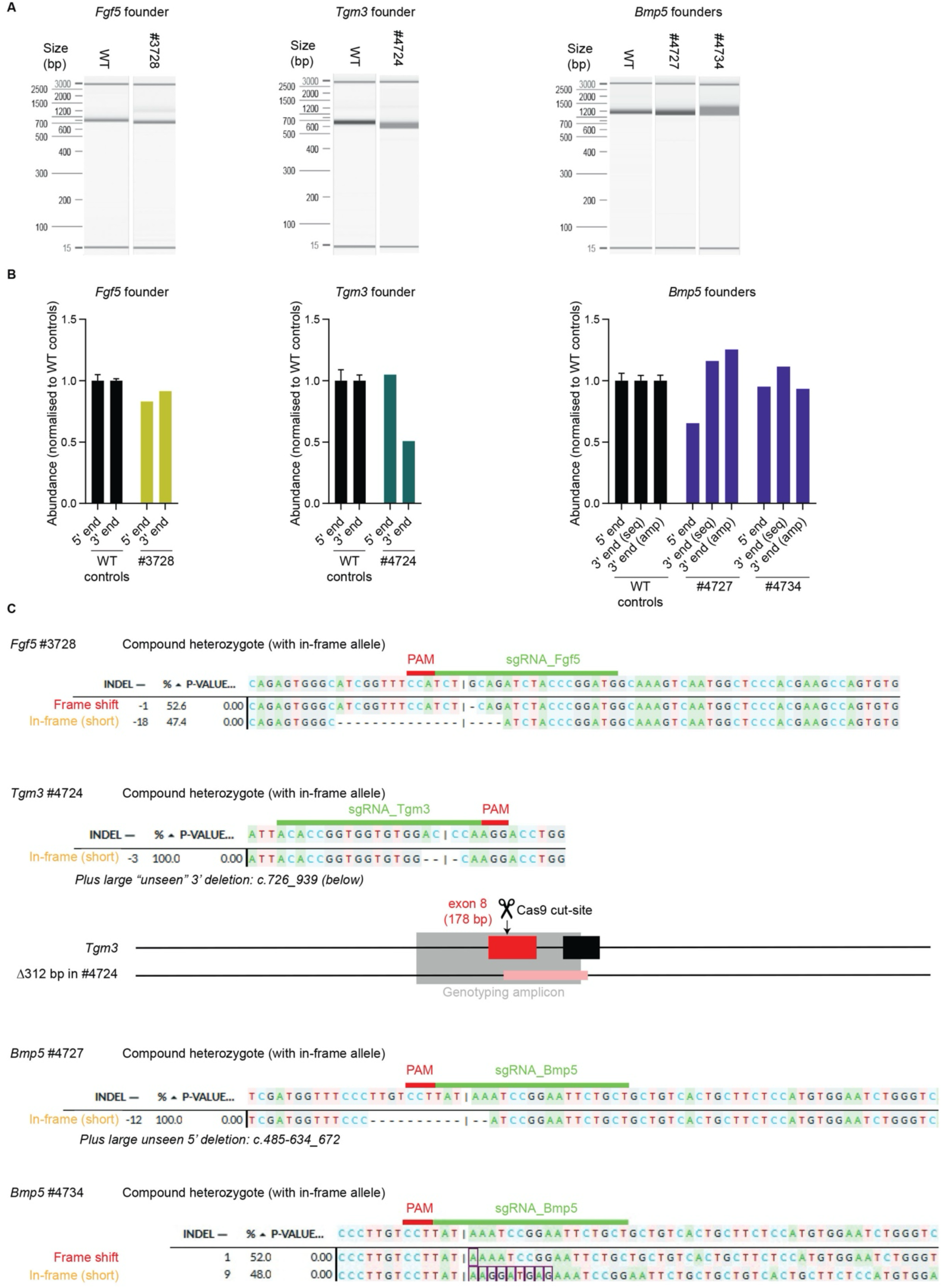
Genotyping analysis of compound heterozygous *Fgf5*, *Tgm3* and *Bmp5* founders carrying one in-frame allele. A. PCR amplification around the targeted site in compound heterozygous founders carrying an in-frame allele: *Fgf5* #3728; *Tgm3* #4724 and *Bmp5* #4727 and #4734. B. qPCR quantification around the 5’ and 3’ genotyping primer annealing sites (and the nested sequencing primer annealing site for *Bmp5*) in these founders, normalised to three wild-type controls (mean ± SEM). C. DECODR deconvolution of Sanger sequence chromatograms of the genotyping amplicon in these founders. A schematic of the genome damage in *Tgm3* founder #2724 (drawn to scale, deleted regions shown in pink). For a schematic of the genome damage in *Bmp5* founder #4727, see Figure 4C. For genotyping analysis of *Tyrp1* founder #4086, see Figures 1 and S1-2; and of *Slc45a2* founders #4062, #4064 and #4079, see Figures 2 and S3.

**Figure S10:**
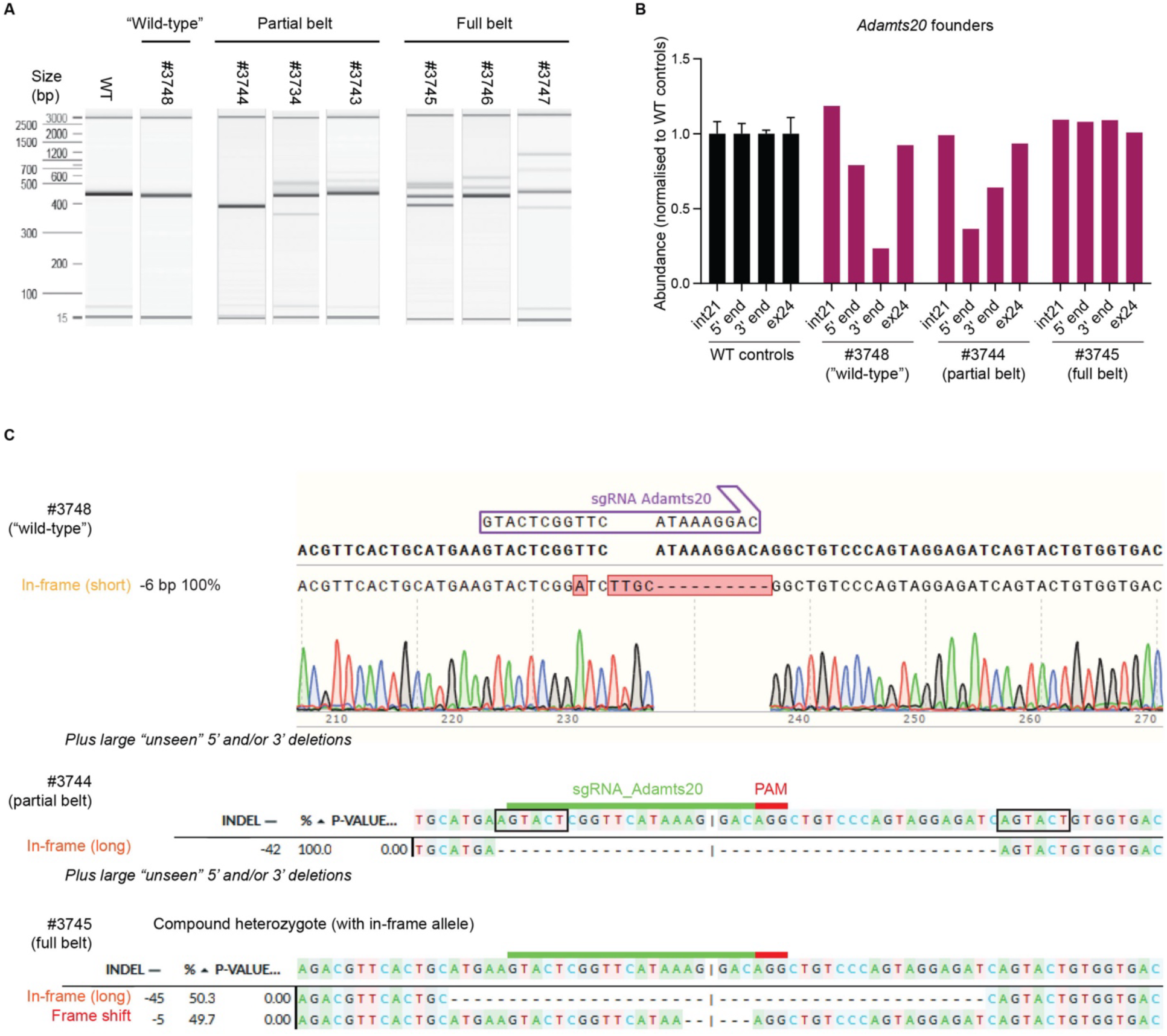
Genotyping analysis of *Adamts20* founders carrying an in-frame allele. A. PCR amplification around the targeted site in *Adamts20* founders carrying an in-frame allele: #3748, #3744 and #3745. B. qPCR quantification around the 5’ and 3’ genotyping primer annealing sites and more distal sites in intron 21 and exon 24 (∼1.5 kb away from the targeted site) in these founders, normalised to three wild-type controls (mean ± SEM). C. Alignment and DECODR deconvolution of Sanger sequence chromatograms of the genotyping amplicon in these founders. The 42 bp deletion in #3744 is flanked by 6 bp of microhomology (boxed).

**Table S1:**
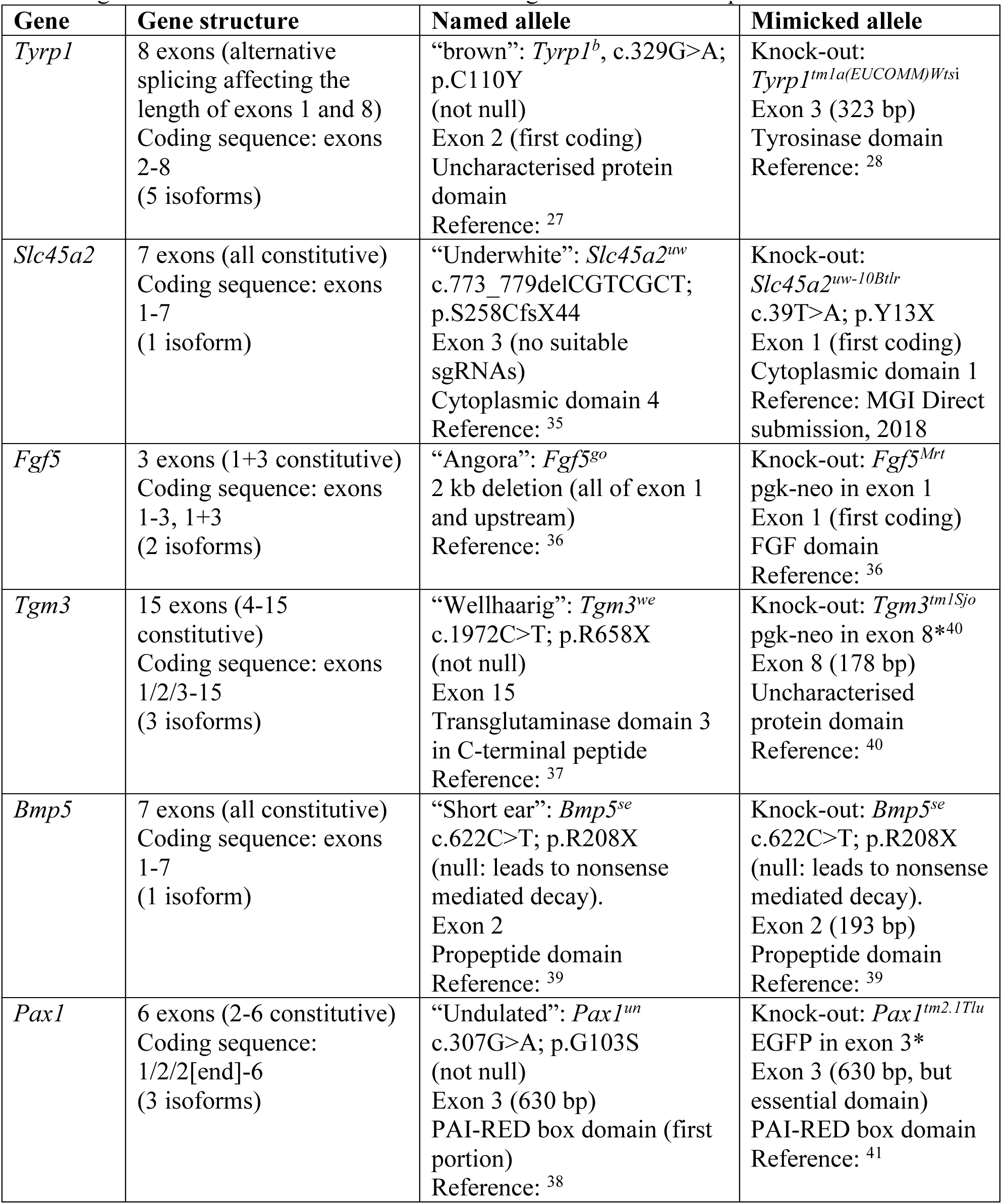
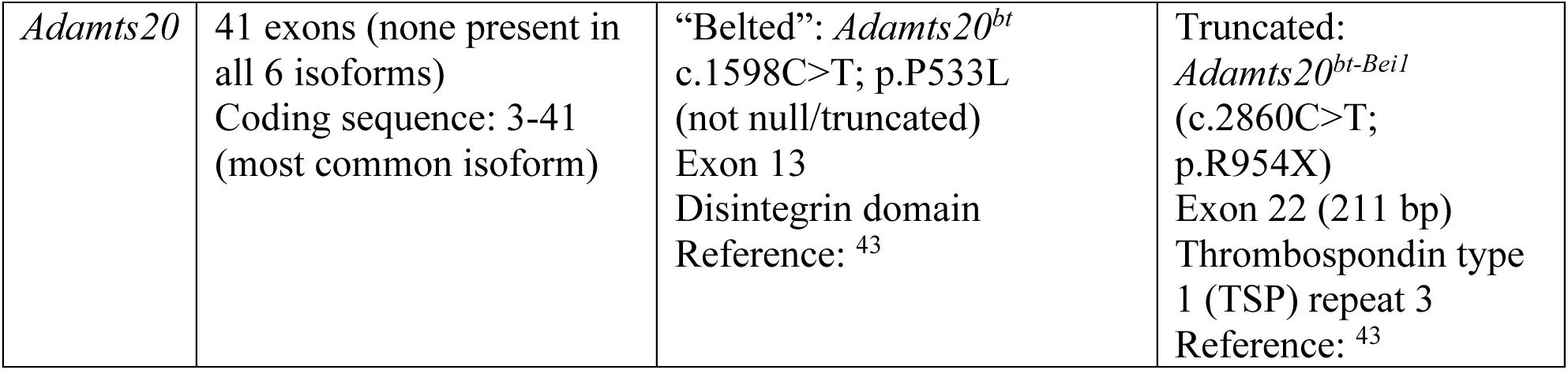
Structure of genes used in this study. *Current gene structure annotation means numbering differs from cited publications.

| Gene | Gene structure | Named allele | Mimicked allele |
| --- | --- | --- | --- |
| *Tyrp1* | 8 exons (alternative splicing affecting the length of exons 1 and 8)<br>Coding sequence: exons 2-8<br>(5 isoforms) | “brown”: *Tyrp1<sup>b</sup>*, c.329G>A; p.C110Y<br>(not null)<br>Exon 2 (first coding)<br>Uncharacterised protein domain<br>Reference: <sup>27</sup> | Knock-out: *Tyrp1<sup>tm1a(EUCOMM)Wtsi</sup>*<br>Exon 3 (323 bp)<br>Tyrosinase domain<br>Reference: <sup>28</sup> |
| *Slc45a2* | 7 exons (all constitutive)<br>Coding sequence: exons 1-7<br>(1 isoform) | “Underwhite”: *Slc45a2<sup>uw</sup>*<br>c.773_779delCGTCGCT; p.S258CfsX44<br>Exon 3 (no suitable sgRNAs)<br>Cytoplasmic domain 4<br>Reference: <sup>35</sup> | Knock-out: *Slc45a2<sup>uw-10Btlr</sup>*<br>c.39T>A; p.Y13X<br>Exon 1 (first coding)<br>Cytoplasmic domain 1<br>Reference: MGI Direct submission, 2018 |
| *Fgf5* | 3 exons (1+3 constitutive)<br>Coding sequence: exons 1-3, 1+3<br>(2 isoforms) | “Angora”: *Fgf5<sup>go</sup>*<br>2 kb deletion (all of exon 1 and upstream)<br>Reference: <sup>36</sup> | Knock-out: *Fgf5<sup>Mrt</sup>*<br>pgk-neo in exon 1<br>Exon 1 (first coding)<br>FGF domain<br>Reference: <sup>36</sup> |
| *Tgm3* | 15 exons (4-15 constitutive)<br>Coding sequence: exons 1/2/3-15<br>(3 isoforms) | “Wellhaarig”: *Tgm3<sup>we</sup>*<br>c.1972C>T; p.R658X<br>(not null)<br>Exon 15<br>Transglutaminase domain 3 in C-terminal peptide<br>Reference: <sup>37</sup> | Knock-out: *Tgm3<sup>tm1Sjo</sup>*<br>pgk-neo in exon 8*<sup>40</sup><br>Exon 8 (178 bp)<br>Uncharacterised protein domain<br>Reference: <sup>40</sup> |
| *Bmp5* | 7 exons (all constitutive)<br>Coding sequence: exons 1-7<br>(1 isoform) | “Short ear”: *Bmp5<sup>se</sup>*<br>c.622C>T; p.R208X<br>(null: leads to nonsense mediated decay).<br>Exon 2<br>Propeptide domain<br>Reference: <sup>39</sup> | Knock-out: *Bmp5<sup>se</sup>*<br>c.622C>T; p.R208X<br>(null: leads to nonsense mediated decay).<br>Exon 2 (193 bp)<br>Propeptide domain<br>Reference: <sup>39</sup> |
| *Pax1* | 6 exons (2-6 constitutive)<br>Coding sequence: 1/2/2[end]-6<br>(3 isoforms) | “Undulated”: *Pax1<sup>un</sup>*<br>c.307G>A; p.G103S<br>(not null)<br>Exon 3 (630 bp)<br>PAI-RED box domain (first portion)<br>Reference: <sup>38</sup> | Knock-out: *Pax1<sup>tm2.1Tlu</sup>*<br>EGFP in exon 3*<br>Exon 3 (630 bp, but essential domain)<br>PAI-RED box domain<br>Reference: <sup>41</sup> |
| *Adamts20* | 41 exons (none present in all 6 isoforms)<br>Coding sequence: 3-41<br>(most common isoform) | “Belted”: *Adamts20<sup>bt</sup>*<br>c.1598C>T; p.P533L<br>(not null/truncated)<br>Exon 13<br>Disintegrin domain<br>Reference: <sup>43</sup> | Truncated:<br>*Adamts20<sup>bt-Beil</sup>*<br>(c.2860C>T;<br>p.R954X)<br>Exon 22 (211 bp)<br>Thrombospondin type<br>1 (TSP) repeat 3<br>Reference: <sup>43</sup> |

**Table S2:** Founders from single guide zygote EP genotyping results. * Founders in photographs. † DECODR results. ‡ Other software

| Gene | ID, sex | Phenotype | % frame shift<br>(no. of products)<br>† | % in-frame<br>(no. of products)<br>† | % total indel<br>† | Notes |
| --- | --- | --- | --- | --- | --- | --- |
| *Tyrp1* | #4083*<br>♂ | Brown | 69.7%<br>(3) | ≥21 bp<br>23.6% (1)<br><21 bp:<br>6.7% (1) | 100% | Plus larger indel(s):<br>results don't reflect<br>PCR product sizes |
| *Tyrp1* | #4084*<br>♂ | Brown | 27.4%<br>(3) | ≥21 bp<br>37.6% (2)<br><21 bp:<br>8.1% (2) | 73.1% | WT = 26.9%;<br>Plus larger indel(s):<br>results don't reflect<br>PCR product sizes;<br>DAS: WT = 10.7%;<br>2 dupA = 0.4% |
| *Tyrp1* | #4085*<br>♂ | Brown | 71.9%<br>(8) | ≥21 bp<br>0% (0)<br><21 bp:<br>28.1% (2) | 100% | Plus larger indel(s):<br>results don't reflect<br>PCR product sizes |
| *Tyrp1* | #4086<br>♀ | Brown | 0%<br>(0) | ≥21 bp<br>0% (0)<br><21 bp:<br>100% (1) | 100% | Δ12 bp (in-frame)<br>c.636_647del<br>p. D212_H215del No<br>additional flanking<br>mutations;<br>Compound<br>heterozygous for<br>above mutation and<br>large deletion<br>spanning 5'<br>genotyping primer<br>annealing site:<br>c.794_933delinsCTT<br>p.I169Lfs*49 |
| *Slc45a2* | #4074*<br>♂ | Underwhite | 100%<br>(2) | ≥21 bp<br>0% (0)<br><21 bp:<br>0% (0) | 100% | Plus larger indel(s):<br>results don't reflect<br>PCR product sizes |
| *Slc45a2* | #4075*<br>♂ | Underwhite | 100%<br>(2) | ≥21 bp<br>0% (0)<br><21 bp:<br>0% (0) | 100% | Plus larger indel(s):<br>results don't reflect<br>PCR product sizes |
| *Slc45a2* | #4069<br>♂ | Patchy | 15.0%<br>(3) | ≥21 bp<br>0% (0)<br><21 bp:<br>85% (1) | 100% | Plus larger indel(s):<br>results don't reflect<br>PCR product sizes |
| *Slc45a2* | #4070<br>♂ | Patchy | Analysis<br>failed | Analysis<br>failed | Analysis<br>failed | Deconvolution<br>analysis failed.<br>Multiple PCFR<br>product sizes. |
| *Slc45a2* | #4071<br>♂ | Patchy | 100%<br>(2) | ≥21 bp<br>0% (0)<br><21 bp:<br>0% (0) | 100% | Plus larger indel(s):<br>results don't reflect<br>PCR product sizes |
| *Slc45a2* | #4072*<br>♂ | Patchy | 73.1%<br>(7) | ≥21 bp<br>26.9% (1)<br><21 bp:<br>0% (0) | 100% | Plus larger indel(s):<br>results don't reflect<br>PCR product sizes |
| *Slc45a2* | #4073*<br>♂ | Patchy | 100%<br>(2) | ≥21 bp<br>0% (0)<br><21 bp:<br>0% (0) | 100% | Plus larger indel(s):<br>results don't reflect<br>PCR product sizes |
| *Slc45a2* | #4080*<br>♀ | Patchy | 100%<br>(3) | ≥21 bp<br>0% (0)<br><21 bp:<br>0% (0) | 100% | Plus larger indel(s):<br>results don't reflect<br>PCR product sizes |
| *Slc45a2* | #4060<br>♂ | Black<br>(WT) | 73.0%<br>(3) | ≥21 bp<br>0% (0)<br><21 bp:<br>27.0% (1) | 100% | Plus larger indel(s):<br>results don't reflect<br>PCR product sizes |
| *Slc45a2* | #4061<br>♂ | Black<br>(WT) | 51.7%<br>(2) | ≥21 bp<br>0% (0)<br><21 bp:<br>48.3% (1) | 100% | Plus larger indel(s):<br>results don't reflect<br>PCR product sizes |
| *Slc45a2* | #4062<br>♂ | Black<br>(WT) | 100%<br>(1) | ≥21 bp<br>0% (0)<br><21 bp:<br>0% (0) | 100% | ‡ (Found by<br>alignment, in-frame)<br>c.36_42delCTATCAA<br>ins[AA,385+647_<br>385+566]<br>p.S2_Q14del |
|  |  |  |  |  |  | No additional flanking mutations;<br>Compound heterozygous for above mutation and large deletion spanning 3' genotyping primer annealing site:<br>c.1-1_385+93del |
| *Slc45a2* | #4063<br>♂ | Black (WT) | 85.5% (1) | ≥21 bp<br>14.6% (1)<br><21 bp:<br>0% (0) | 100% | Plus larger indel(s):<br>results don't reflect PCR product sizes |
| *Slc45a2* | #4064<br>♂ | Black (WT) | 50.5% (1) | ≥21 bp<br>0% (0)<br><21 bp:<br>49.5% (1) | 100% | Compound heterozygote:<br>Δ9 bp (in-frame)<br>c.35_43del<br>CCTATCAAT<br>p.T12T, Y13_S15del;<br>Δ7 bp (FS)<br>c.34_40del<br>ACCTATC<br>p.T12NfsX3 |
| *Slc45a2* | #4065<br>♂ | Black (WT) | 84.5% (4) | ≥21 bp<br>0% (0)<br><21 bp:<br>15.5% (1) | 100% | Plus larger indel(s):<br>results don't reflect PCR product sizes |
| *Slc45a2* | #4066<br>♂ | Black (WT) | 53.8% (2) | ≥21 bp<br>0% (0)<br><21 bp:<br>46.2% (1) | 100% | Plus larger indel(s):<br>results don't reflect PCR product sizes |
| *Slc45a2* | #4067<br>♂ | Black (WT) | 51.6% (1) | ≥21 bp<br>0% (0)<br><21 bp:<br>48.4% (1) | 100% | Plus larger indel(s):<br>results don't reflect PCR product sizes |
| *Slc45a2* | #4068<br>♂ | Black (WT) | 48.3% (1) | ≥21 bp<br>51.7% (1)<br><21 bp:<br>0% (0) | 100% | Plus larger indel(s):<br>results don't reflect PCR product sizes |
| *Slc45a2* | #4076*<br>♀ | Black (WT) | 49.2% (1) | ≥21 bp<br>0% (0)<br><21 bp:<br>50.8% (1) | 100% | Plus larger indel(s):<br>results don't reflect PCR product sizes |
| *Slc45a2* | #4077*<br>♀ | Black<br>(WT) | 100%<br>(4) | ≥21 bp<br>0% (0)<br><21 bp:<br>0% (0) | 100% | Plus larger indel(s):<br>results don't reflect<br>PCR product sizes |
| *Slc45a2* | #4078*<br>♀ | Black<br>(WT) | Analysis<br>failed | Analysis<br>failed | Analysis<br>failed | Deconvolution<br>analysis failed.<br>Multiple band sizes. |
| *Slc45a2* | #4079*<br>♀ | Black<br>(WT) | 50.8%<br>(1) | ≥21 bp<br>0% (0)<br><21 bp:<br>49.3% (1) | 100% | Compound<br>heterozygote:<br>Δ6 bp (in-frame)<br>c.32_37delATACCT<br>p.H11H, T12_Y13del<br>(inDelphi: 8.4%);<br>Δ4 bp (FS)<br>c.33_36delTACC<br>p.H11HfsX5<br>(inDelphi: 13.8%) |
| *Fgf5* | #3712<br>♂ | Angora | 38.1%<br>(1) | ≥21 bp<br>50.4% (1)<br><21 bp:<br>11.5% (1) | 100% |  |
| *Fgf5* | #3713<br>♂ | Angora | 100%<br>(1) | ≥21 bp<br>0% (0)<br><21 bp:<br>0% (0) | 100% | Δ1 bp (FS)<br>c.298del<br>p.Q100RfsX17<br>(inDelphi: 3.2%);<br>Compound<br>heterozygous for<br>above mutation and<br>large deletion<br>spanning 3'<br>genotyping primer<br>annealing site |
| *Fgf5* | #3716<br>♂ | Angora | 100%<br>(2) | ≥21 bp<br>0% (0)<br><21 bp:<br>0% (0) | 100% |  |
| *Fgf5* | #3720<br>♂ | Angora | 100%<br>(2) | ≥21 bp<br>0% (0)<br><21 bp:<br>0% (0) | 100% |  |
| *Fgf5* | #3723*<br>♂ | Angora | Analysis<br>failed | Analysis<br>failed | Analysis<br>failed | Deconvolution<br>analysis failed.<br>Multiple PCR product<br>sizes. |
| *Fgf5* | #3724<br>♂ | Angora | 100%<br>(2) | ≥21 bp<br>0% (0) | 100% | Main (95.8%, Δ4 bp,<br>FS): |
|  |  |  |  | <21 bp:<br>0% (0) |  | c. 295_298del<br>CTGC<br>p.L99RfsX17;<br>Almost entirely<br>compound<br>heterozygous for<br>above mutation and<br>large deletion<br>spanning 5'<br>genotyping primer<br>annealing site |
| *Fgf5* | #3727<br>♂ | Angora | 100%<br>(1) | ≥21 bp<br>0% (0)<br><21 bp:<br>0% (0) | 100% | Plus larger indel(s):<br>results don't reflect<br>PCR product sizes |
| *Fgf5* | #3728<br>♀ | Angora | 52.6%<br>(1) | ≥21 bp<br>0% (0)<br><21 bp:<br>47.4% (1) | 100% | Compound<br>heterozygote:<br>Δ18 bp (in-frame):<br>c.283_300del<br>p.I95_Q100del;<br>Δ1 bp (FS):<br>c.279del<br>p.L99LfsX18 |
| *Fgf5* | #3730<br>♀ | Angora | 100%<br>(2) | ≥21 bp<br>0% (0)<br><21 bp:<br>0% (0) | 100% | Plus larger indel(s):<br>results don't reflect<br>PCR product sizes |
| *Fgf5* | #3733<br>♀ | Angora | 100%<br>(3) | ≥21 bp<br>0% (0)<br><21 bp:<br>0% (0) | 100% | Plus larger indel(s):<br>results don't reflect<br>PCR product sizes |
| *Tgm3* | #3735<br>♂ | Wellhaarig | 100%<br>(1) | ≥21 bp<br>0% (0)<br><21 bp:<br>0% (0) | 100% | Δ8 bp (FS):<br>c.739_746del<br>p.P247LfsX4<br>high MH product<br>(inDelphi: 44.5%);<br>Compound<br>heterozygous for<br>above mutation and<br>large deletion<br>spanning 5'<br>genotyping primer<br>annealing site |
| *Tgm3* | #3736<br>♂ | Wellhaarig | 100%<br>(1) | ≥21 bp<br>0% (0) | 100% | Δ8 bp (FS):<br>c.739_746del<br>p.P247LfsX4 |
|  |  |  |  | <21 bp:<br>0% (0) |  | high MH product<br>(inDelphi: 44.5%);<br>Compound<br>heterozygous for<br>above mutation and<br>large deletion<br>spanning 5' and 3'<br>genotyping primer<br>annealing sites |
| *Tgm3* | #3737*<br>♂ | Wellhaarig | 100%<br>(1) | ≥21 bp<br>0% (0)<br><21 bp:<br>0% (0) | 100% | -250 bp (FS, extends<br>into intron):<br>c.670-178_742delinsC<br>‡ (Found by<br>alignment);<br>Compound<br>heterozygous for<br>above mutation and<br>large deletion<br>spanning 5'<br>genotyping primer<br>annealing site |
| *Tgm3* | #3738<br>♀ | Wellhaarig | 100%<br>(1) | ≥21 bp<br>0% (0)<br><21 bp:<br>0% (0) | 100% | Δ8 bp (FS):<br>c.739-746del<br>p.P247LfsX4<br>high MH product<br>(inDelphi: 44.5%);<br>Compound<br>heterozygous for<br>above mutation and<br>large deletion<br>spanning 3'<br>genotyping primer<br>annealing site |
| *Tgm3* | #3739<br>♀ | Wellhaarig | 100%<br>(2) | ≥21 bp<br>0% (0)<br><21 bp:<br>0% (0) | 100% | Plus larger indel(s):<br>results don't reflect<br>PCR product sizes |
| *Tgm3* | #3740<br>♀ | Wellhaarig | 100%<br>(2) | ≥21 bp<br>0% (0)<br><21 bp:<br>0% (0) | 100% | High MH product =<br>64.9%;<br>Plus larger indel(s):<br>results don't reflect<br>PCR product sizes |
| *Tgm3* | #3741<br>♀ | Wellhaarig | 79.4%<br>(2) | ≥21 bp<br>0% (0)<br><21 bp:<br>20.6% (1) | 100% | Plus larger indel(s):<br>results don't reflect<br>PCR product sizes |
| *Tgm3* | #3742<br>♀ | Wellhaarig | 100%<br>(2) | ≥21 bp<br>0% (0)<br><21 bp:<br>0% (0) | 100% | Plus larger indel(s):<br>results don't reflect<br>PCR product sizes |
| *Bmp5* | #3714<br>♂ | Short ear | 98%<br>(2) | ≥21 bp<br>0% (0)<br><21 bp:<br>0% (0) | 98% | Main (84.3%)<br>c. 601_602insT<br>p.Y201LfsX2<br>(inDelphi: 14.1%)<br>WT = 2%;<br>(In spleen, WT = 0%) |
| *Bmp5* | #3715<br>♂ | Short ear | 96.4%<br>(1) | ≥21 bp<br>0% (0)<br><21 bp:<br>3.6% (1) | 100% | Main (96.4%, Δ2 bp,<br>FS):<br>c.600_601del<br>p.I200IfsX2<br>(inDelphi = 5.2%);<br>Almost entirely<br>compound<br>heterozygous for<br>above mutation and<br>large deletion<br>spanning 5'<br>genotyping primer<br>annealing site |
| *Bmp5* | #3717<br>♂ | Short ear | 100%<br>(2) | ≥21 bp<br>0% (0)<br><21 bp:<br>0% (0) | 100% | Plus larger indel(s):<br>results don't reflect<br>PCR product sizes |
| *Bmp5* | #3718<br>♂ | Short ear | 100%<br>(3) | ≥21 bp<br>0% (0)<br><21 bp:<br>0% (0) | 100% | Plus larger indel(s):<br>results don't reflect<br>PCR product sizes |
| *Bmp5* | #3719<br>♂ | Short ear | 100%<br>(2) | ≥21 bp<br>0% (0)<br><21 bp:<br>0% (0) | 100% | Compound<br>heterozygote:<br>Δ38 bp (FS):<br>c.583_620del<br>p.A195SfsX3;<br>Δ28 bp (FS):<br>c.583_610del<br>p.A195RfsX32 |
| *Bmp5* | #3721*<br>♂ | Short ear | 100%<br>(3) | ≥21 bp<br>0% (0)<br><21 bp:<br>0% (0) | 100% | Plus larger indel(s):<br>results don't reflect<br>PCR product sizes |
| *Bmp5* | #3722<br>♂ | Short ear | (1) | ≥21 bp<br>0%<br>(0) | 100% | ‡ (Found by alignment<br>after gel extraction of<br>each band) |
|  |  |  |  | <21 bp:<br>(1) |  | Δ12,ins2 bp (FS):<br>c. 595_606delinsTG,<br>p.R199WfsX34;<br>Δ498 bp (spanning<br>exon 2 + intron 2):<br>c.554_683+368del;<br>Plus large deletion(s)<br>spanning 5' and 3'<br>genotyping primer<br>annealing sites |
| *Bmp5* | #3725<br>♀ | Short ear;<br>Grey<br>patches | 100%<br>(1) | ≥21 bp<br>0% (0)<br><21 bp:<br>0% (0) | 100% | Δ10 bp (FS):<br>c.599_608del<br>p.I200TfsX33<br>(inDelphi = 14.2%);<br>Mosaic for above<br>mutation and large<br>deletions spanning 5'<br>genotyping primer<br>annealing site, some<br>extending to *Myo5a*<br>("dilute") locus |
| *Bmp5* | #3726<br>♀ | Short ear | 66.7%<br>(5) | ≥21 bp<br>15.2% (1)<br><21 bp:<br>0% (0) | 81.9% | WT = 18.1%<br>Plus larger indel(s):<br>results don't reflect<br>PCR product sizes;<br>‡ WT = 4% by ICE;<br>(In spleen, WT = 0%) |
| *Bmp5* | #3729<br>♀ | Short ear | 76.0%<br>(3) | ≥21 bp<br>24.0%<br>(2)<br><21 bp:<br>0% (0) | 100% | Plus larger indel(s):<br>results don't reflect<br>PCR product sizes |
| *Bmp5* | #3731<br>♀ | Short ear | 98.6%<br>(3) | ≥21 bp<br>0% (0)<br><21 bp:<br>0% (0) | 98.6% | WT = 1.4%<br>Plus larger indel(s):<br>results don't reflect<br>PCR product sizes;<br>(In spleen, WT = 0%) |
| *Bmp5* | #3732<br>♀ | Short ear | 100%<br>(3) | ≥21 bp<br>0% (0)<br><21 bp:<br>0% (0) | 100% | Plus larger indel(s):<br>results don't reflect<br>PCR product sizes |
| *Pax1* | #4081<br>♂ | Undulated | 100%<br>(1) | ≥21 bp<br>0% (0)<br><21 bp:<br>0% (0) | 100% | Δ4 bp (FS):<br>c.339_343del<br>p.R113RfsX12<br>(inDelphi: 21.7%); |
|  |  |  |  |  |  | Compound heterozygous for above mutation and large deletion spanning 3' genotyping primer annealing site |
| *Pax1* | #4082*<br>♂ | Undulated | 100%<br>(4) | ≥21 bp<br>0% (0)<br><21 bp:<br>0% (0) | 100% | Plus larger indel(s): results don't reflect PCR product sizes |
| *Pax1* | #4087<br>♀ | Undulated | 92.9%<br>(4) | ≥21 bp<br>0% (0)<br><21 bp:<br>7.1% (1) | 100% | Plus large deletion(s) spanning 5' and 3' genotyping primer annealing sites |
| *Pax1* | #4088<br>♀ | Undulated | 100%<br>(3) | ≥21 bp<br>0% (0)<br><21 bp:<br>0% (0) | 100% | All Δ 8 bp<br>‡ ICE result (DECODR only detects most abundant product);<br>Plus large deletion spanning 5' genotyping primer annealing sites |
| *Pax1* | #4089<br>♀ | Undulated;<br>Humane endpoint at 3 weeks | 100%<br>(4) | ≥21 bp<br>0% (0)<br><21 bp:<br>0% (0) | 100% | Plus larger indel(s): results don't reflect PCR product sizes |

**Table S3:** Mutagenesis is consistent across all three germ layers. † DECODR results. ‡ Other software.

| Gene | ID, sex | Tail tip (largely ectoderm)† | Spleen (mesoderm)† | Liver (endoderm)† |
| --- | --- | --- | --- | --- |
| *Pax1* | #4081<br>♂ | c.339_343del<br>p.R113RfsX12<br>(inDelphi: 21.7%) | 100% FS (2 products):<br>Δ5bp: 91.8%<br>c.339_343del<br>p.R113RfsX12<br>(inDelphi: 21.7%)<br>Δ4 bp: 8.3% | c.339_343del<br>p.R113RfsX12<br>(inDelphi: 21.7%) |
| *Pax1* | #4082<br>♂ | 100% FS (4 products):<br>Δ277 bp: 50.3%,<br>Δ98 bp: 39.4%,<br>Δ5 bp: 6.7%,<br>Δ4 bp: 3.6% | 100% FS (4 products):<br>Δ277 bp: 28.7%,<br>Δ98 bp: 51.3%,<br>Δ5 bp: 13.4%,<br>Δ4 bp: 6.6% | 100% FS (4 products):<br>Δ277 bp: 19.6%,<br>Δ98 bp: 56.5%,<br>Δ5 bp: 15.6%,<br>Δ4 bp: 8.3% |
|  |  | Plus larger indel(s):<br>results don't reflect Qx<br>sizes | Plus larger indel(s):<br>results don't reflect Qx<br>sizes | Plus larger indel(s):<br>results don't reflect Qx<br>sizes |
| *Pax1* | #4087<br>♀ | 92.7% FS (4 products):<br>Δ5 bp: 61.8%<br>c.339_343del<br>p.R113RfsX12<br>(inDelphi: 21.7%)<br>Δ4 bp: 20.7%,<br>Δ41 bp: 5.4%,<br>Δ19 bp: 5.0%<br>7.1% In-frame (1<br>product):<br>Δ18 bp: 7.1% | c.339_343del<br>p.R113RfsX12<br>(inDelphi: 21.7%) | 100% FS (2 products):<br>Δ5bp: 89.4%<br>c.339_343del<br>p.R113RfsX12<br>(inDelphi: 21.7%)<br>Δ4 bp: 10.6% |
| *Pax1* | #4088<br>♀ | 100% FS (3 products):<br>Δ8a bp: 60%<br>(inDelphi: 7.5%),<br>Δ8b bp: 39%,<br>Δ8c bp: 1%<br>(‡ ICE result) | 100% FS (2 products):<br>Δ8a bp: 67%<br>(inDelphi: 7.5%),<br>Δ8b bp: 33%,<br>(‡ ICE result) | 100% FS (2 products):<br>Δ8a bp: 62%<br>(inDelphi: 7.5%),<br>Δ8b bp: 38%,<br>(‡ ICE result) |
| *Pax1* | #4089<br>♀ | 100% FS (4 products):<br>Δ74 bp: 62.1%,<br>Δ14a bp: 21.8%,<br>Δ97a bp: 10.5%,<br>Δ13 bp: 5.6%,<br>Plus larger indel(s):<br>results don't reflect Qx<br>sizes | 94.3% FS (5 products):<br>Δ74 bp: 48.5%,<br>Δ14b bp: 24.3%,<br>Δ97b bp: 17.5%,<br>Δ47 bp: 4.0%,<br>5.7% In-frame (1<br>product):<br>Δ6 bp: 5.7%<br>Plus larger indel(s):<br>results don't reflect Qx<br>sizes | 100% FS (4 products):<br>Δ74 bp: 57.1%,<br>Δ14b bp: 25.8%,<br>Δ97b bp: 10.9%,<br>Δ13 bp: 6.3%,<br>Plus larger indel(s):<br>results don't reflect Qx<br>sizes |

**Table S4:**
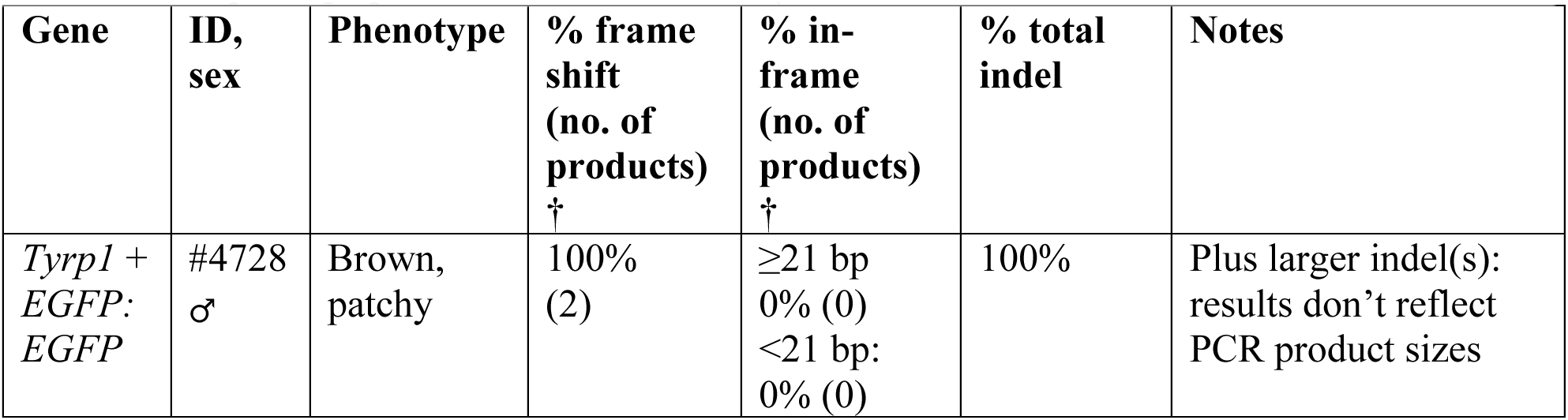
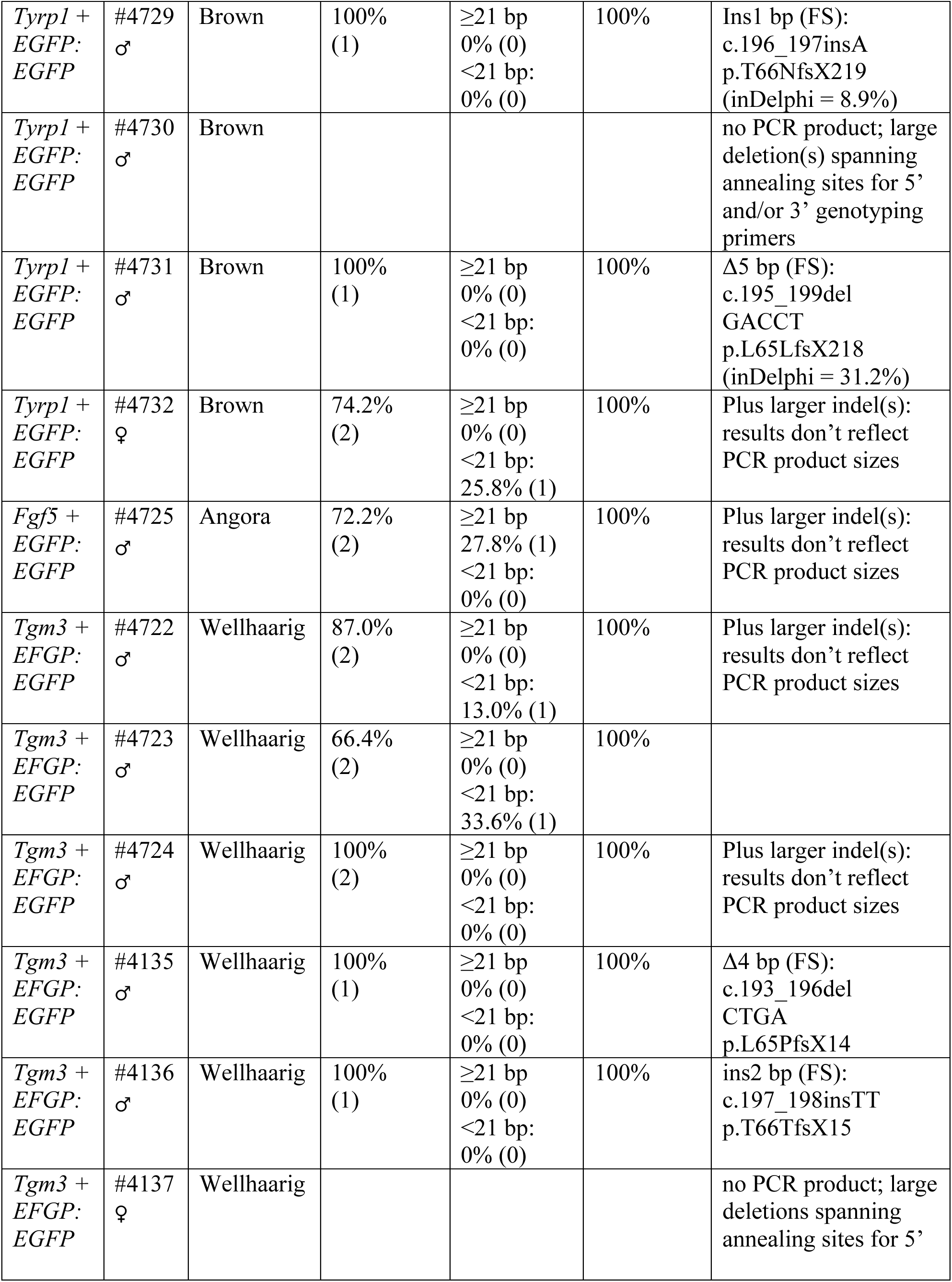
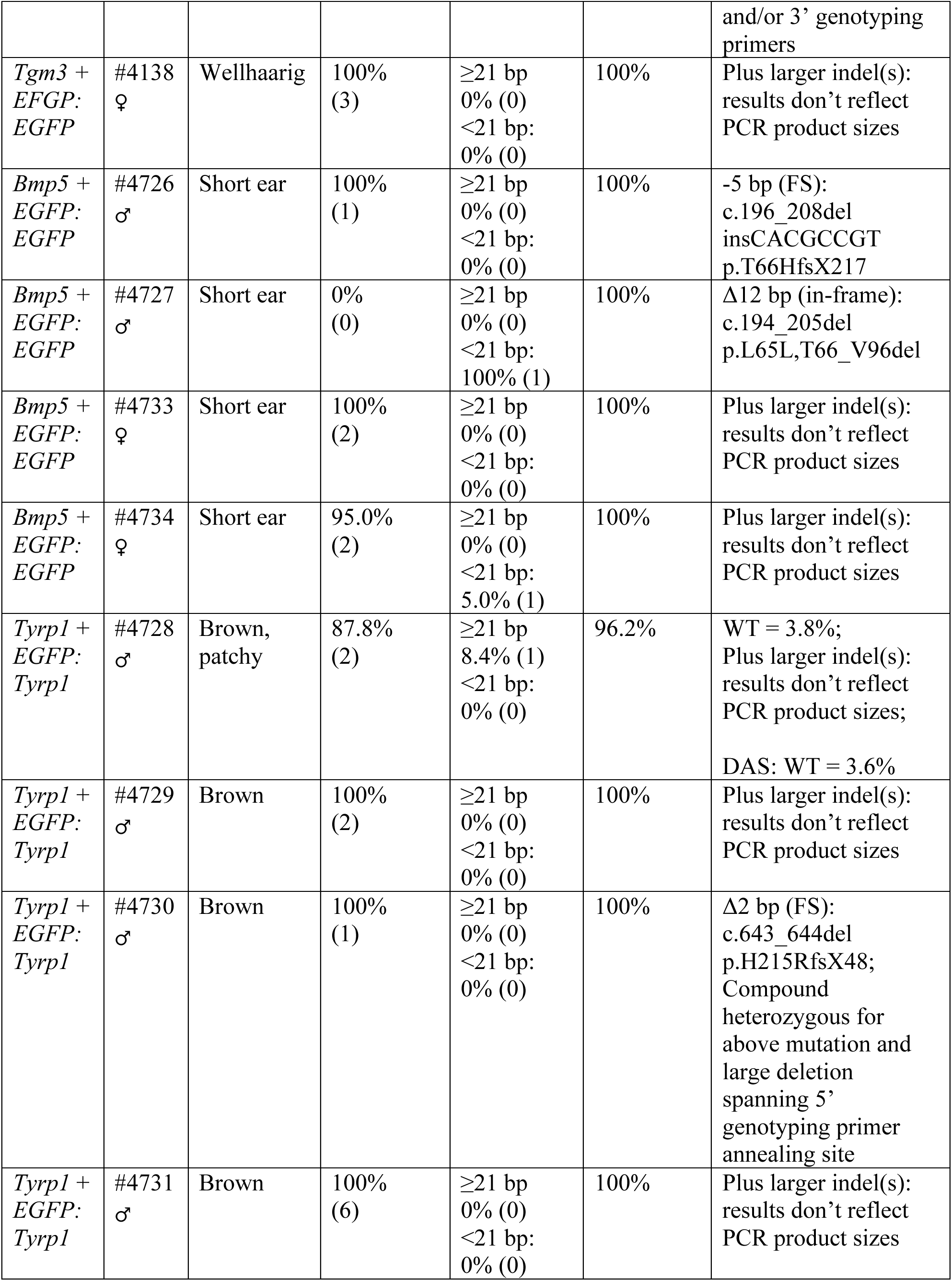
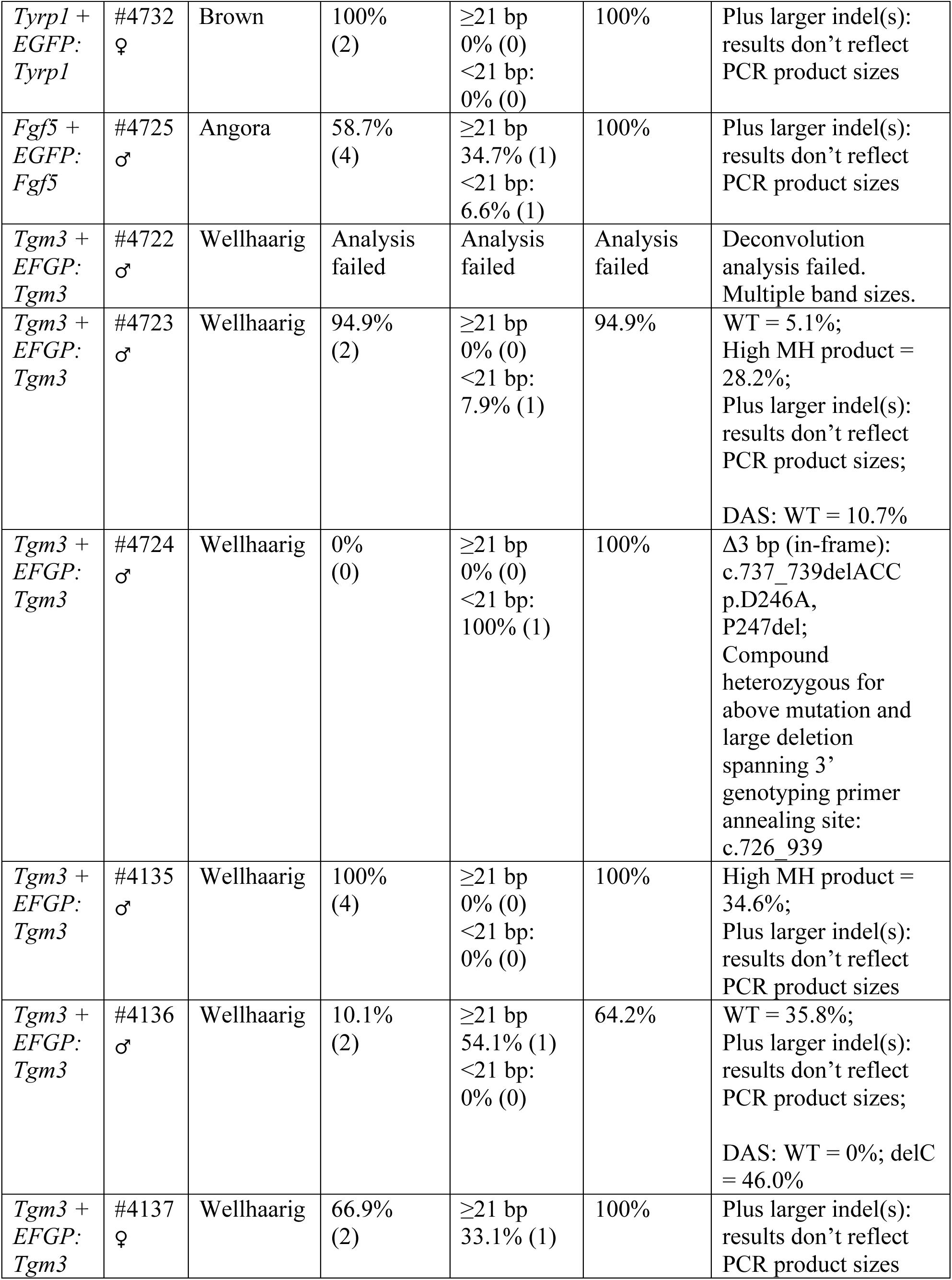
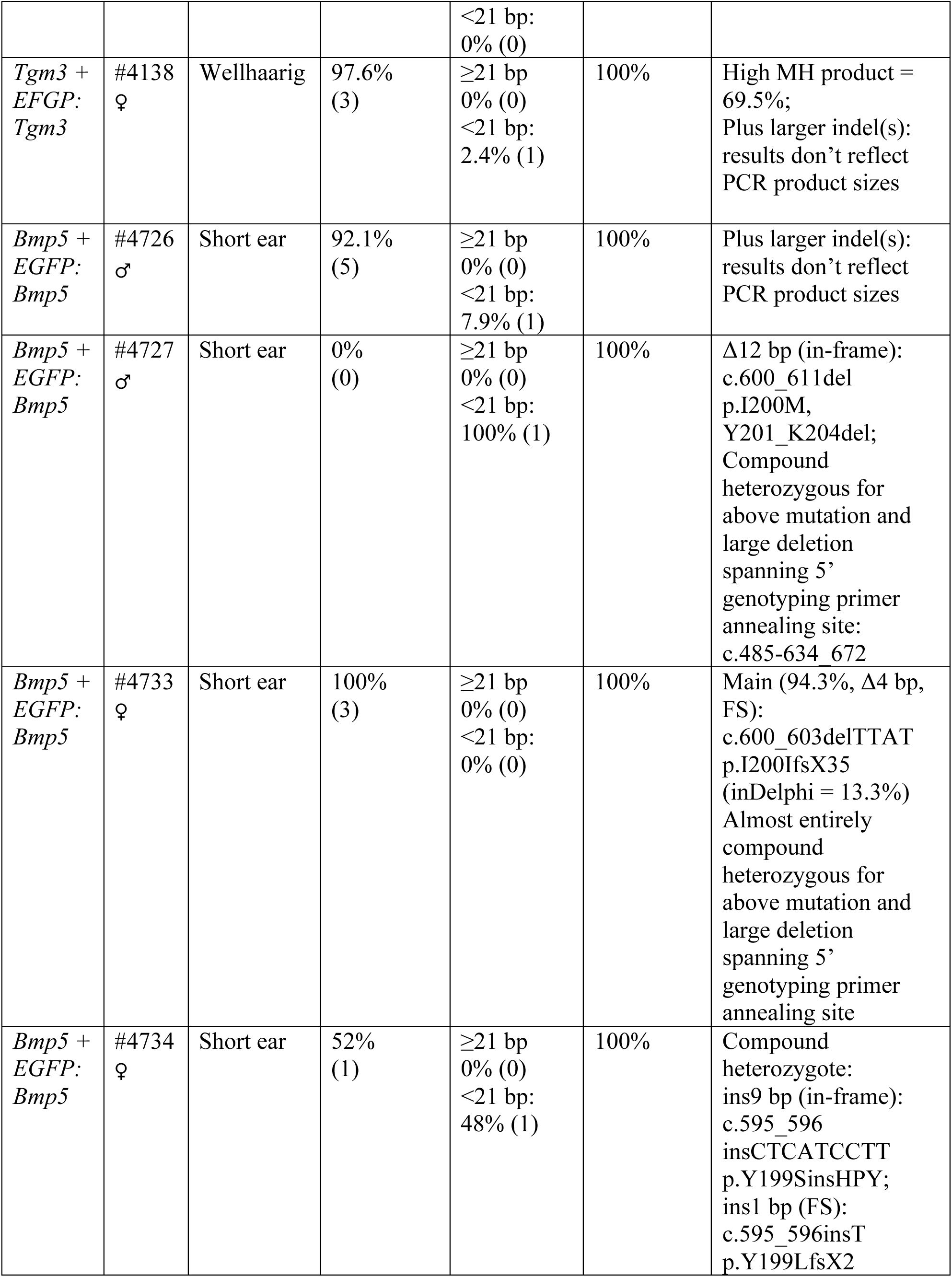
Founders from two guides (endogenous gene + EGFP) zygote EP genotyping results. * Founders in photographs. † DECODR results. ‡ Other software

**Table S5:** Founders from two guides (two endogenous genes) zygote EP genotyping results. Founders in photographs. † DECODR results. ‡ Other software

| Gene | ID, sex | Phenotype | % frame shift<br>(no. of products)<br>† | % in-frame<br>(no. of products)<br>† | % total indel | Notes |
| --- | --- | --- | --- | --- | --- | --- |
| *Tyrp1* + *Bmp5*:<br>*Tyrp1* | #5486*<br>♀ | Brown + Short ear | 33.4%<br>(3) | ≥21 bp<br>22.3% (1)<br><21 bp:<br>0% (0) | 55.6% | WT = 44.3%;<br>Plus larger indel(s): results don't reflect PCR product sizes;<br>DAS: WT = 0.6%;<br>dupA = 30.8% |
| *Tyrp1* + *Tgm3*:<br>*Tyrp1* | #5485*<br>♀ | Brown + Wellhaarig | 100%<br>(2) | ≥21 bp<br>0% (0)<br><21 bp:<br>0% (0) | 100% | Plus larger indel(s): results don't reflect PCR product sizes |
| *Tyrp1* + *Bmp5*:<br>*Bmp5* | #5486*<br>♀ | Brown + Short ear | 100%<br>(3) | ≥21 bp<br>0% (0)<br><21 bp:<br>0% (0) | 100% | Plus larger indel(s): results don't reflect PCR product sizes |
| *Tyrp1* + *Tgm3*:<br>*Tgm3* | #5485*<br>♀ | Brown + Wellhaarig | 100%<br>(2) | ≥21 bp<br>0% (0)<br><21 bp:<br>0% (0) | 100% | High MH product = 49.2%;<br>Plus larger indel(s): results don't reflect PCR product sizes |
| *Tgm3* + *Adamts20*:<br>*Tgm3* | #2983*<br>♂ | Wellhaarig + Full belt | 100%<br>(2) | ≥21 bp<br>0% (0)<br><21 bp:<br>0% (0) | 100% | High MH product = 47.2%;<br>Plus larger indel(s): results don't reflect PCR product sizes |
| *Tgm3* + *Adamts20*:<br>*Tgm3* | #2984*<br>♂ | Wellhaarig + Full belt |  |  |  | Very faint PCR products; large deletion(s) spanning annealing sites for 5' and/or 3' genotyping primers |
| *Tgm3* + *Adamts20*:<br>*Tgm3* | #2985*<br>♀ | Wellhaarig + Full belt | 100%<br>(2) | ≥21 bp<br>0% (0)<br><21 bp:<br>0% (0) | 100% | High MH product = 27.2%;<br>Plus larger indel(s): results don't reflect PCR product sizes |
| *Tgm3* + *Adamts20*: *Tgm3* | #2986*<br>♀ | Wellhaarig + Full belt | 100% (1) | ≥21 bp<br>0% (0)<br><21 bp:<br>0% (0) | 100% | Plus larger indel(s): results don't reflect PCR product sizes |
| *Tgm3* + *Adamts20*: *Adamts20* | #2983*<br>♂ | Wellhaarig + Full belt | 100% (1) | ≥21 bp<br>0% (0)<br><21 bp:<br>0% (0) | 100% | Δ17 bp (FS):<br>c.2798_2814del<br>p.H933PfsX9;<br>Compound heterozygous for above mutation and large deletion spanning 5' and 3' genotyping primer annealing sites |
| *Tgm3* + *Adamts20*: *Adamts20* | #2984*<br>♂ | Wellhaarig + Full belt | 100% (1) | ≥21 bp<br>0% (0)<br><21 bp:<br>0% (0) | 100% | Ins1 bp (FS):<br>c.2804_2805insG<br>p.G935GfsX13<br>(inDelphi = 3.8%);<br>Compound heterozygous for above mutation and large deletion spanning 5' genotyping primer annealing site:<br>c.2709-1173_2709-1158,2709-524_2810 |
| *Tgm3* + *Adamts20*: *Adamts20* | #2985*<br>♀ | Wellhaarig + Full belt | 28.9% (1) | ≥21 bp<br>0% (0)<br><21 bp:<br>71.1% (1) | 100% | Plus larger indel(s): results don't reflect PCR product sizes |
| *Tgm3* + *Adamts20*: *Adamts20* | #2986*<br>♀ | Wellhaarig + Full belt | 100% (1) | ≥21 bp<br>0% (0)<br><21 bp:<br>0% (0) | 100% | ‡ (Found by alignment)<br>+1 bp (FS):<br>c.2803delGinsAA<br>p.G935KfsX13;<br>Compound heterozygous for above mutation and large deletion spanning 5' and 3' genotyping primer annealing sites:<br>c.2709-630_2919+95 |

**Table S6:** Founders from *Adamts20* zygote EP genotyping results. Founders in photographs. † DECODR results. ‡ Other software

| Gene | ID, sex | Phenotype | % frame shift<br>(no. of products)<br>† | % in-frame<br>(no. of products)<br>† | % total indel | Notes |
| --- | --- | --- | --- | --- | --- | --- |
| *Adamst20* | #3745*<br>♀ | Full belt | 49.7%<br>(1) | ≥21 bp<br>50.3% (1)<br><21 bp:<br>0% (0) | 100% | Compound heterozygote:<br>Δ45 bp (long in-frame)<br>c.2782_2826del<br>p.M928_D942del;<br>Δ5 bp (FS)<br>c.2804_2808del<br>GACAG<br>p.K934KfsX12<br>(inDelphi = 16.4%) |
| *Adamst20* | #3746<br>♀ | Full belt | 100%<br>(2) | ≥21 bp<br>0% (0)<br><21 bp:<br>0% (0) | 100% | Plus larger indel(s):<br>results don't reflect<br>PCR product sizes |
| *Adamst20* | #3747<br>♂ | Full belt | 78.7%<br>(3) | ≥21 bp<br>21.3% (1)<br><21 bp:<br>0% (0) | 100% | Plus larger indel(s):<br>results don't reflect<br>PCR product sizes |
| *Adamst20* | #3734<br>♂ | Partial belt | 44.4%<br>(4) | ≥21 bp<br>22.5% (1)<br><21 bp:<br>33.1% (1) | 100% | Plus larger indel(s):<br>results don't reflect<br>PCR product sizes |
| *Adamst20* | #3743*<br>♀ | Partial belt | 69%<br>(2) | ≥21 bp<br>0% (0)<br><21 bp:<br>28% (1) | 97% | ‡ (ICE analysis)<br>Plus larger indel(s):<br>results don't reflect<br>PCR product sizes |
| *Adamst20* | #3744*<br>♀ | Partial belt | 0%<br>(0) | ≥21 bp<br>100% (1)<br><21 bp: 0<br>(0) | 100% | Δ42 bp (long in-frame)<br>c.2792_2833del<br>p.S931C,<br>V932_C945del;<br>Plus large unseen<br>5' - 3' deletions |
| *Adamts20* | #3748<br>♀ | Black<br>(WT) | 0%<br>(0) | ≥21 bp<br>0% (0)<br><21 bp:<br>100% (1) | 100% | -6 bp (in-frame)<br>‡ (Found by<br>alignment)<br>c.2795T>A,<br>2798_2807del<br>insTTGC<br>p.V932_336del<br>insDLA<br>Plus large unseen<br>5' - 3' deletions. |

**Table S7:**
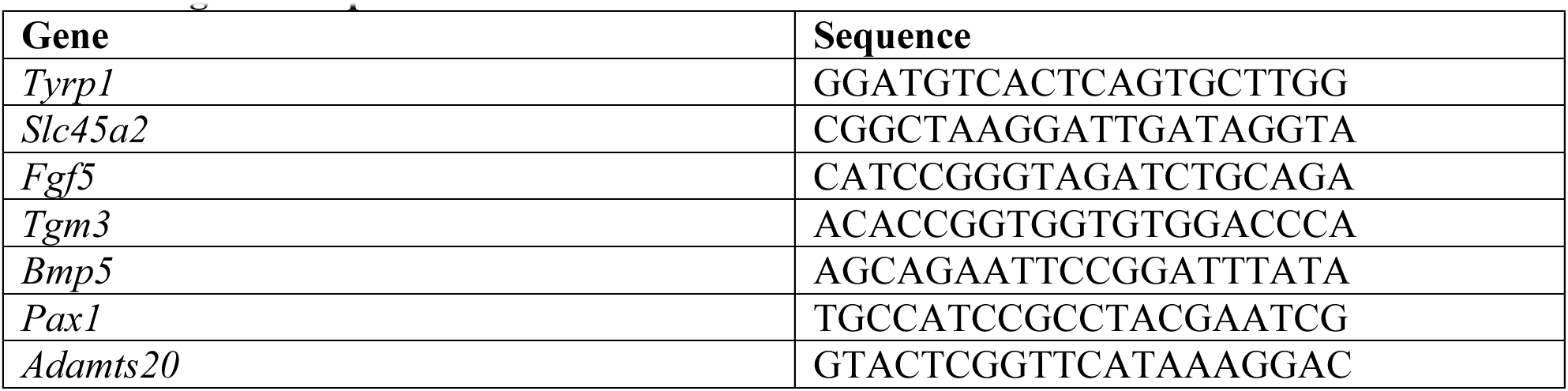
sgRNA sequences.

**Table S8:**
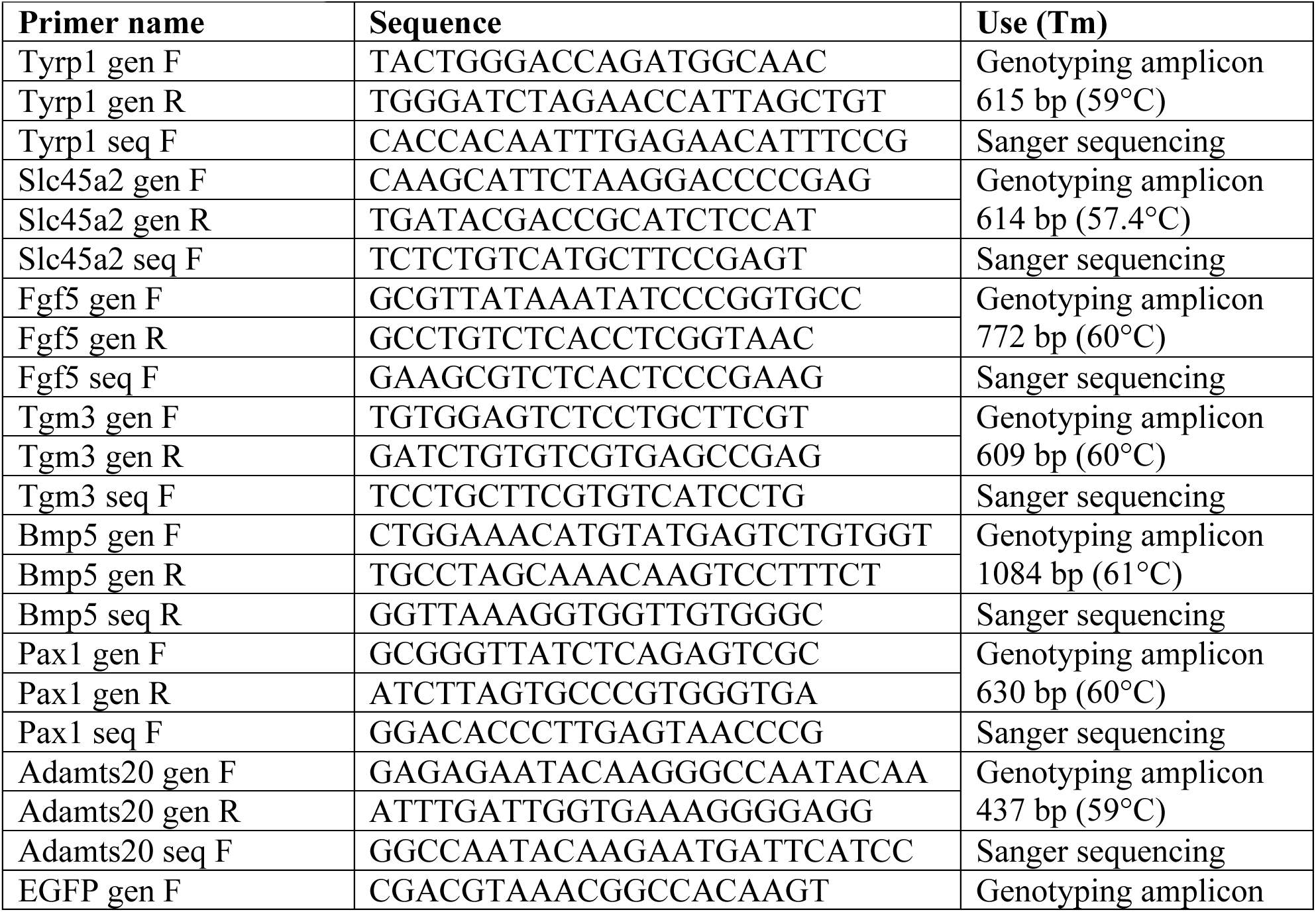
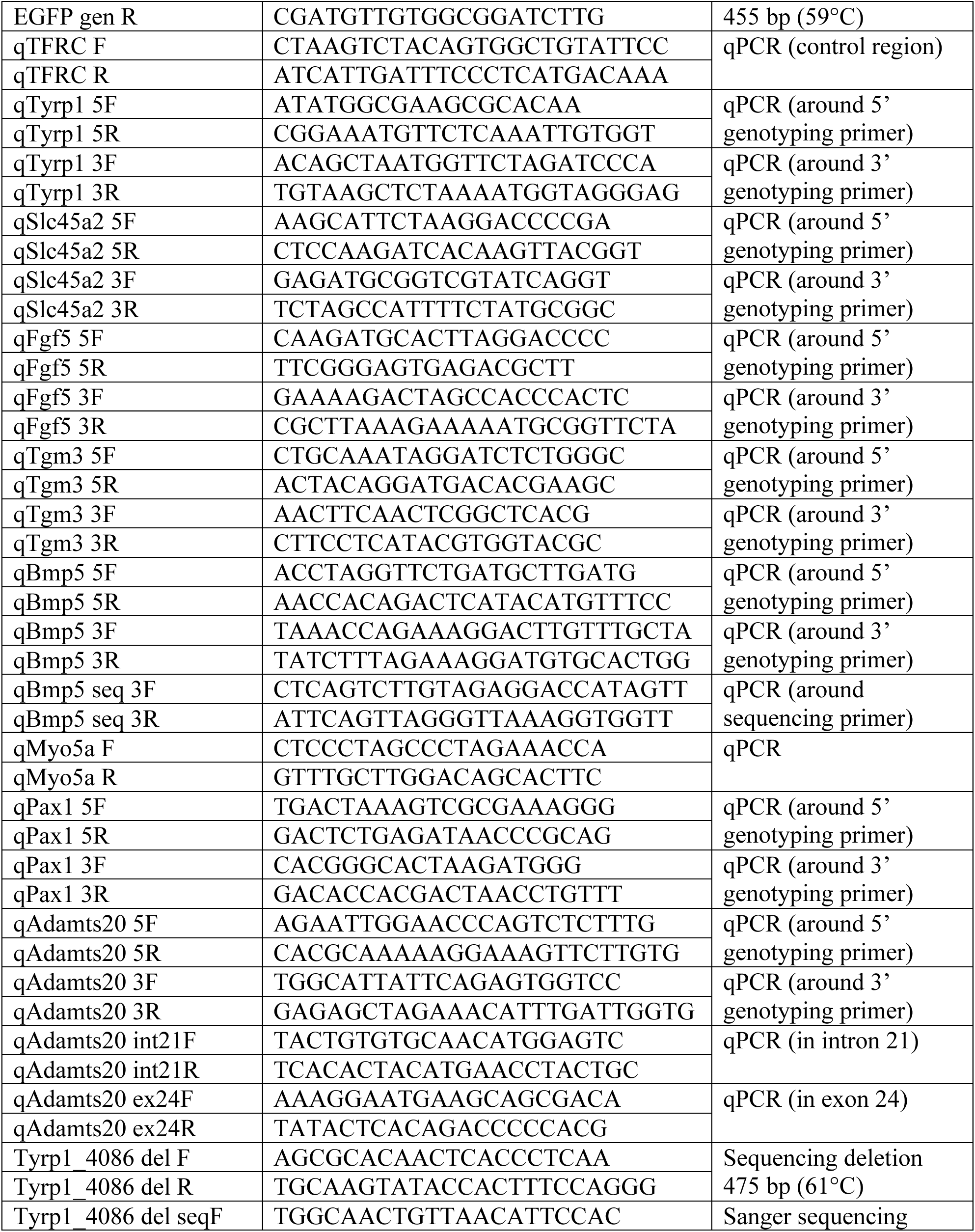
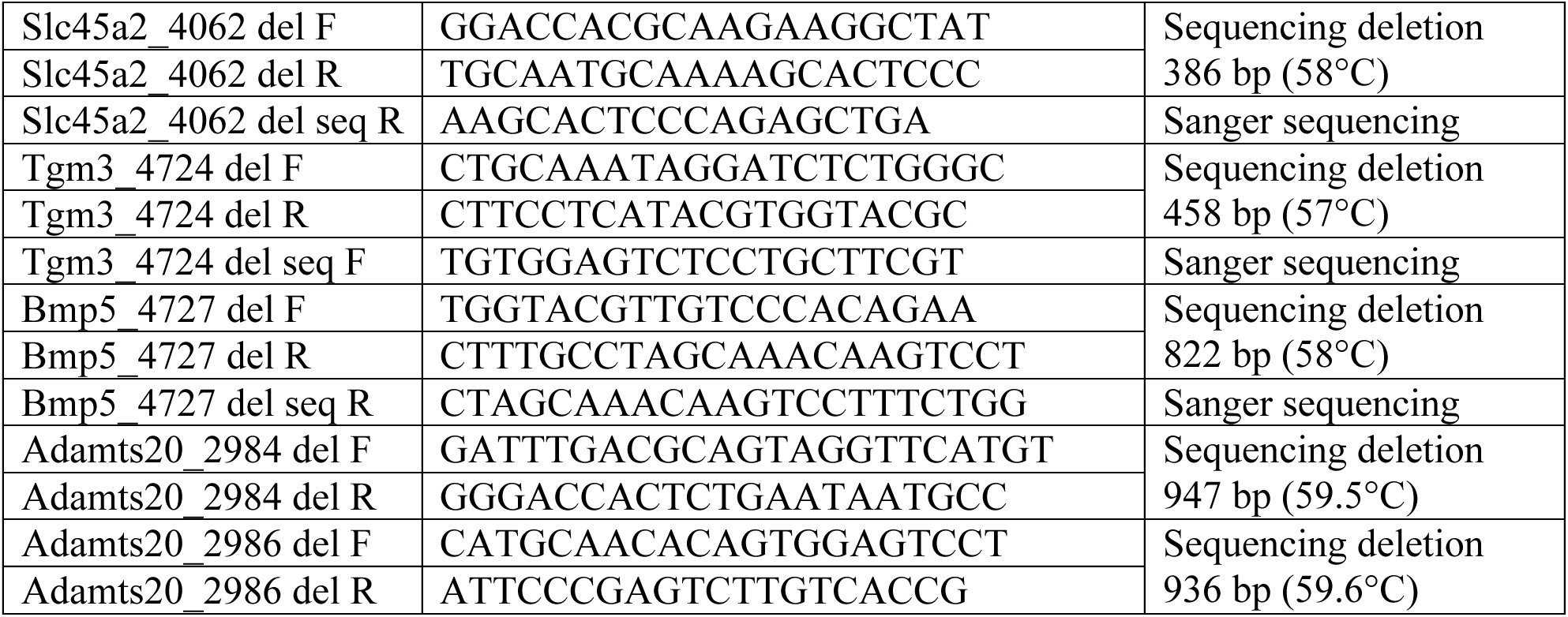
Primers.

**Table S9:**
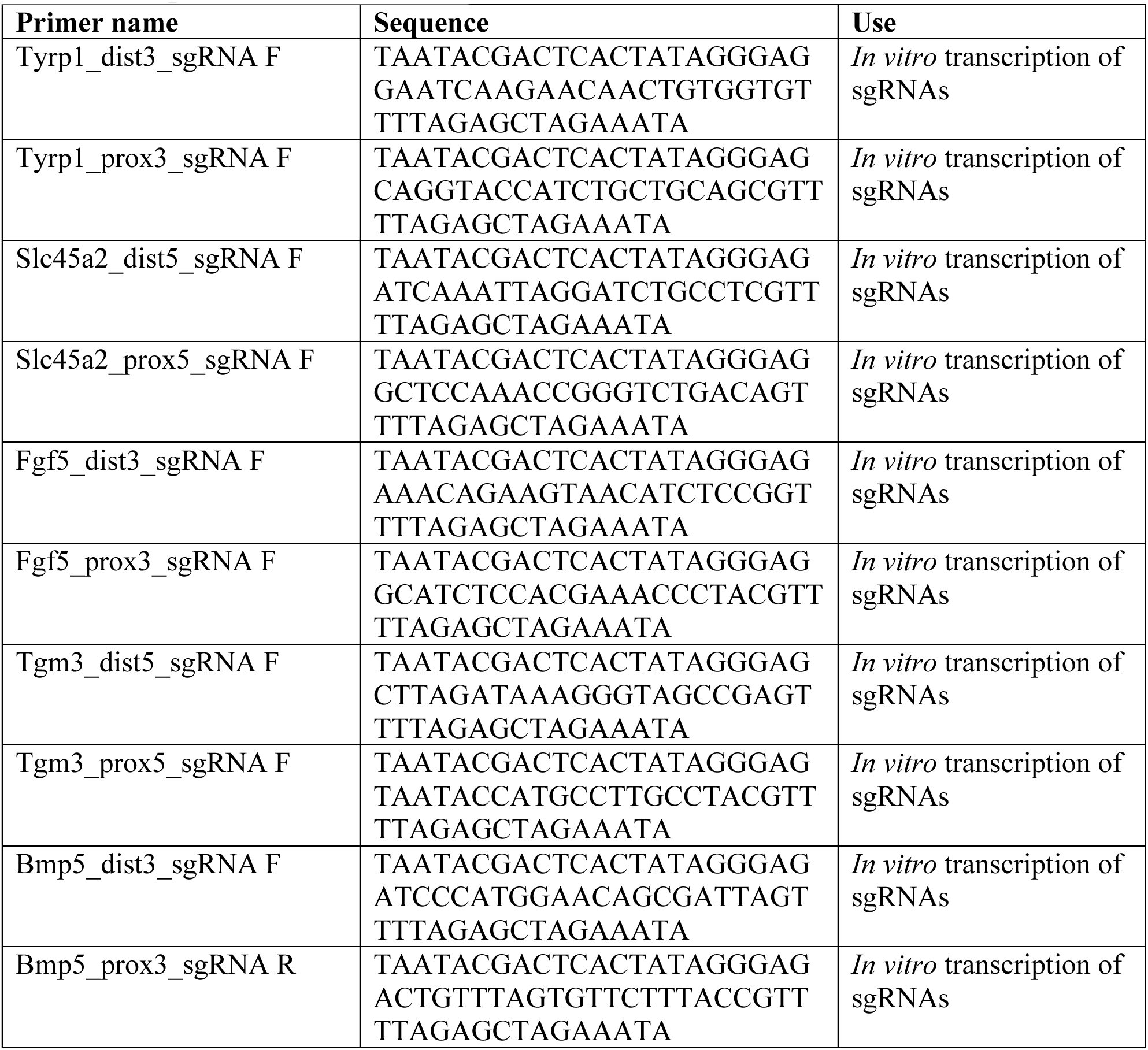
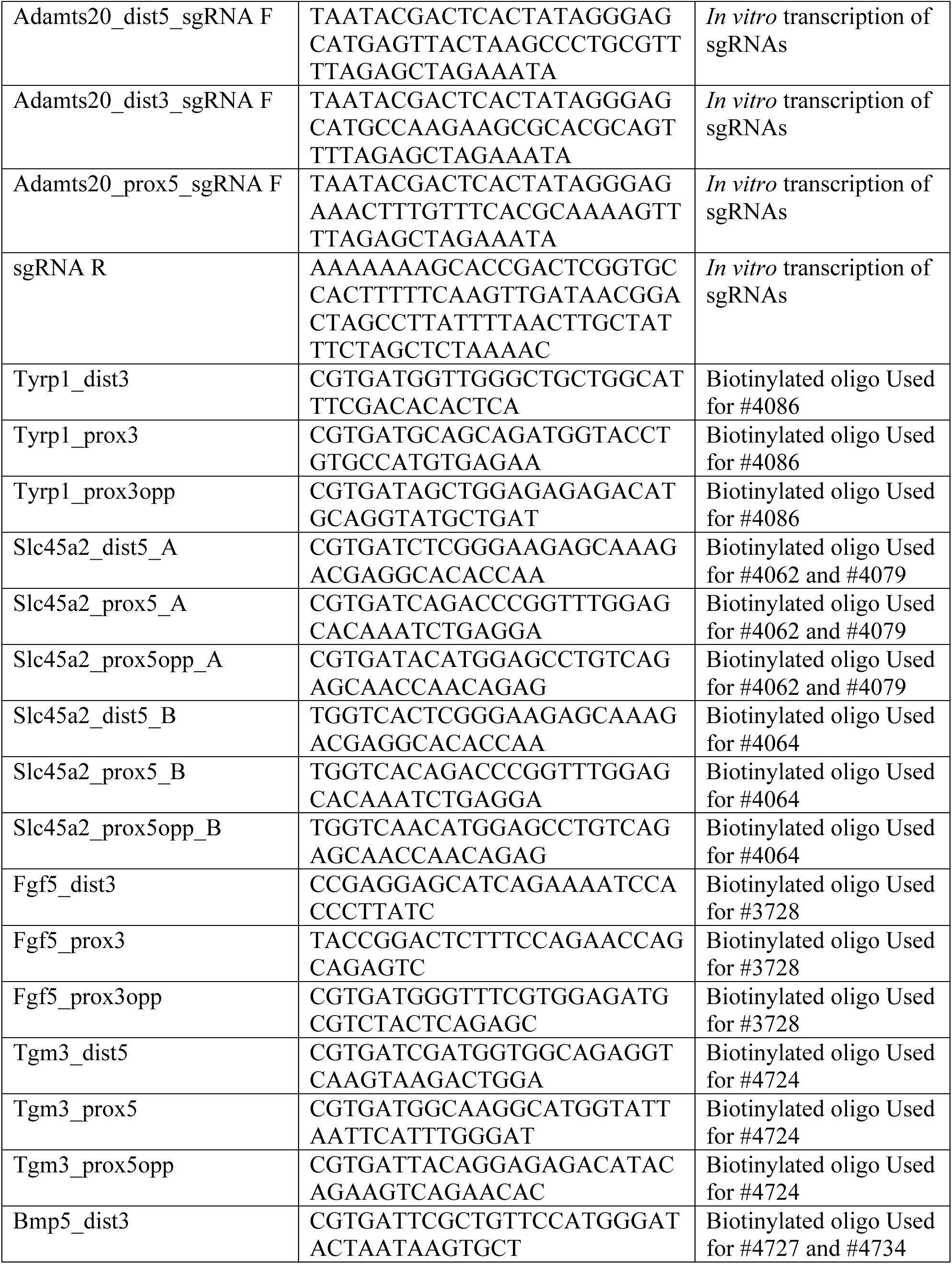
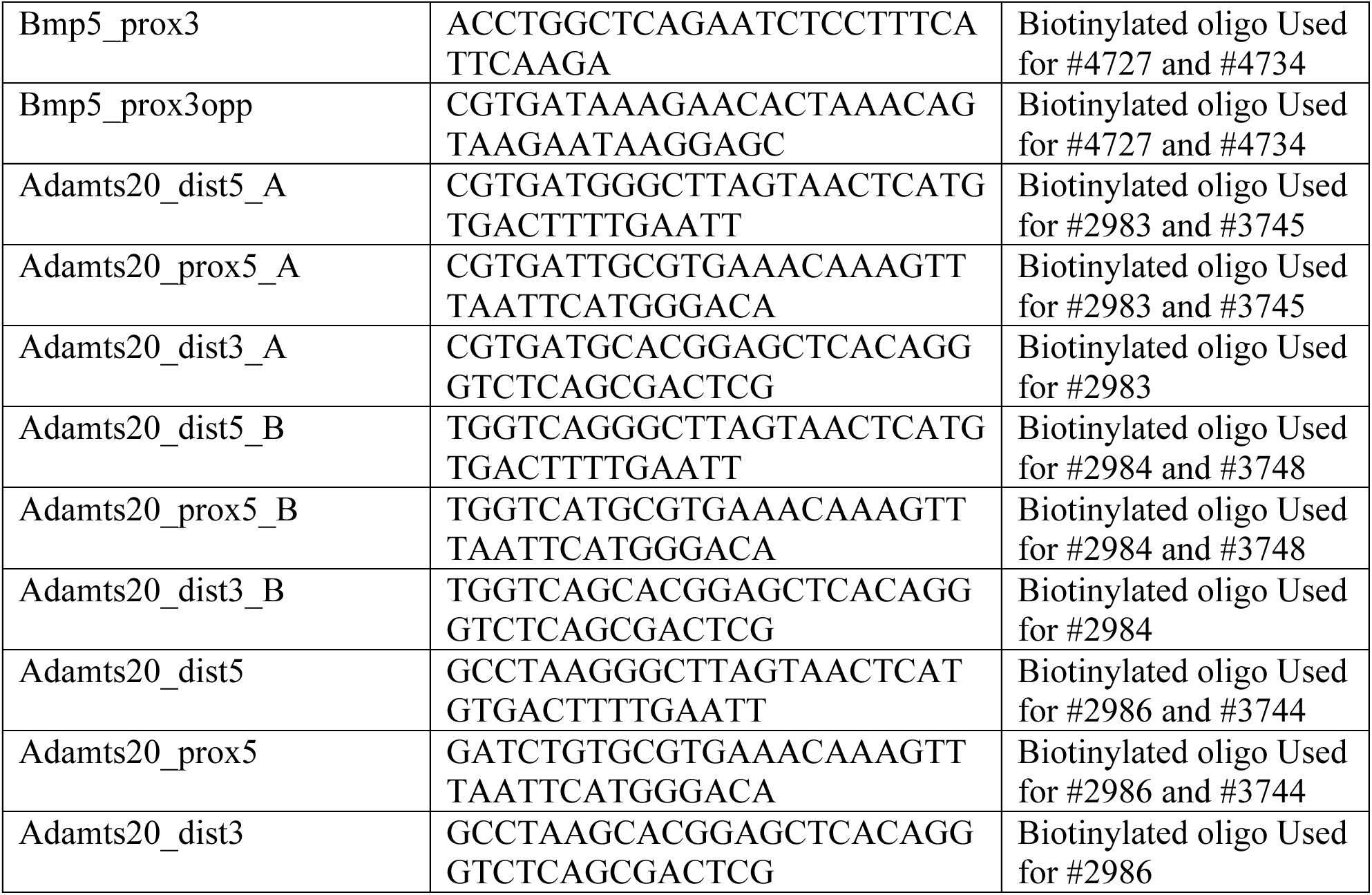
Oligos for ELF-CLAMP.

